# Design principles of dose-response alignment in coupled GTPase switches

**DOI:** 10.1101/2022.06.14.496184

**Authors:** Lingxia Qiao, Pradipta Ghosh, Padmini Rangamani

## Abstract

“Dose-response alignment” (DoRA), where the downstream response of cellular signaling pathways closely matches the fraction of activated receptor, can improve the fidelity of dose information transmission. It is believed that a key component for DoRA is negative feedback and thus a natural question that arises is whether there exist design principles for signaling motifs within such negative feedback loops, which may enable these motifs to attain near-perfect DoRA. Here, we investigated several model formulations of an experimentally validated circuit that couples two molecular switches—mGTPase (monomeric GTPase) and tGTPase (heterotrimeric GTPases) — with negative feedback loops. We find that, in the absence of feedback, the low and intermediate mGTPase activation levels benefit DoRA in the mass action and Hill-function models, respectively. In other cases, where the mass action model with a high mGTPase activation level or the Hill-function model with a non-intermediate mGTPase activation level, the DoRA can be improved by adding negative feedback loops. Furthermore, we found that DoRA in a longer cascade (i.e., tGTPase) can be obtained using Hill-function kinetics under certain conditions. In summary, we show how ranges of activity of mGTPase, reaction kinetics, the negative feedback, and the cascade length affect DoRA. This work provides a framework for improving the DoRA performance in signaling motifs with negative feedback loops.

**Significance Statement:** Dose-response alignment helps cells faithfully transmit dose information; how this alignment is achieved in motifs with negative feedback is unclear. Through rigorous studies interrogating a naturally occurring motif comprised of two species of GTPases coupled by negative feedback loops, this work reveals the versatile roles of negative feedback loops and GTPase regulators on DoRA. We find that the negative feedback can enhance DoRA only with specific kinetic forms and with certain ranges of GTPases activation levels. This knowledge advances our understanding of the role of negative feedback on DoRA and sheds light on the importance of dynamic range of signaling processes as an essential determinant of how cells transfer information about stimuli. Findings can help design signaling circuits with better DoRA behavior, and ultimately augment cell signaling studies.

## 1 Introduction

One of the fundamental challenges in biology is understanding how cells reliably transmit chemical information from the extracellular milieu to the intracellular environment. The classical pathways begin with ligand-receptor interactions and involve complex biochemical reactions on the plasma membrane (1–5). To capture the information transfer for these pathways, several metrics have been proposed. These metrics include dose response alignment (henceforth referred to as DoRA) (4, 6–10), the variance of downstream response (11–16), and the channel capacity of signaling pathways (17–21). Of these metrics, many of them focus on the stochastic behavior of downstream response while the assessment of the DoRA only requires the deterministic response for different levels of the stimulus. DoRA refers to the situation where the dose-response curves of the receptor occupancy and the downstream responses can be closely aligned. For a system exhibiting DoRA, it is thought that the downstream species responds proportionately to the receptor occupancy and thus avoids the amplification of noise caused by hypersensitive response (4, 6). The pervasiveness of DoRA across many systems, implies that it is a trait that is selected during the evolution of regulatory systems and is believed to enhance the fidelity of information transmission (4).

DoRA has been found in many signaling systems, including the *Saccharomyces cerevisiae* pheromone pathway(4), the insulin (22), the thyrotropin (23), angiotensin II (24), and epidermal growth factor (EGF) (25, 26) response systems. Although the biochemical details of these systems are different, the idea that certain network topologies promote DoRA is appealing from the standpoint of identifying design principles. For example, studies have shown that the presence of a negative feedback loop is critical for DoRA (4, 8, 9): the negative feedback loop can increase the level of stimulus that leads to half-maximal activation, which might be the reason for the achievement of DoRA. However, the negative feedback loop alone may not be able to achieve this alignment, and comparator adjusters are required (9). A comparator adjuster can measure the difference between the downstream response and the receptor, and then adjust the downstream response to make it align with the receptor. Such an adjustment is analogous to “proportional control” in engineered systems, where the strength of negative feedback is adjusted to be proportional to the difference between output and input. Besides, a non-feedback topology—”push-pull” topology can also produce perfect DoRA, which was identified in (9) and deeply dissected for coupled molecular switches in (7). For this topology, the downstream response is not only upregulated (push) by the active receptors but also downregulated (pull) by the inactive receptors (7, 9, 27). This mechanism has been validated in *Saccharomyces cerevisiae* yeast cell, where the protein RGS — GTPase activating protein (GAP) protein works as the “pull” by accelerating GTP hydrolysis and thereby, terminating tGTPase signaling (27).

Experimental analyses have revealed that there are several canonical and non-canonical signal transduction pathways through which ligand or stimulus information from the extracellular space is transmitted to the intracellular space and leads to cellular decision making (28–32). Central to these pathways are the large family of GTPase switches, which can switch between GTP-bound (active) state and GDP-bound (inactive) state. The switch from the inactive to the active state is catalyzed by guanine exchange factors (GEFs), and the reverse reaction GTPase activating proteins (GAPs). Recent research used the simplified network topology to identify principles of perfect DoRA for GTPase switches (7, 9). However, these works neglect the effects of GEFs and GAPs, and lack detailed analyses for those circuits occurring frequently in nature but without the identified principle of perfect DoRA. More importantly, signal transduction in cells has more than one switch in series and coupling between switches must be considered.

Due to the ubiquitousness of the DoRA (22–26), a natural question is whether DoRA can be identified in larger signal transduction paradigms. Specifically, can coupled GTPase switches achieve DoRA for intracellular signal transmission? To answer this question, we studied how the DoRA can be improved for an experimentally validated circuit—coupled mGTPase (monomeric GTPase) and tGTPase (heterotrimeric GTPases) switches with negative feedback loops (33) (Figure 1A). GEFs and GAPs are considered in this network, and the core motif is the negative feedback, which is prevalent in signaling pathways (34–38). We investigated several model formulations of this coupled network including the circuit without or with the negative feedback, the choice of model equations, and the logic for two species co-regulating the same target (Figure 1B-C). Through theoretical analyses and numerical simulations, we identified several DoRA design principles including the role of feedback and the length of the cascade in attaining perfect DoRA.

**Figure 1:**
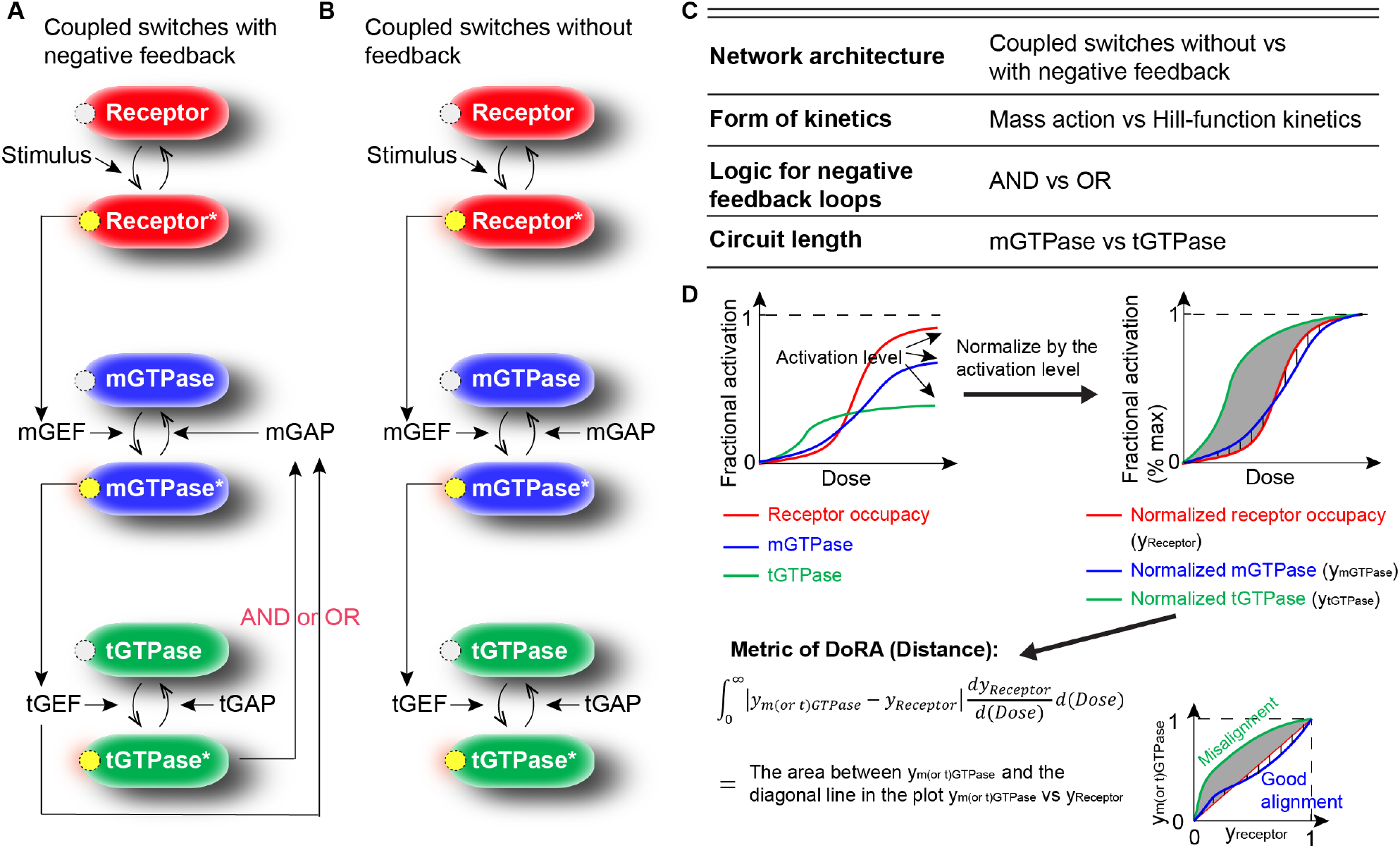
Exploring mechanisms for dose-response alignment (DoRA) through GTPases switches. (A-B) Circuits for coupled switches with negative feedback (A) or without feedback (B). For the circuit in (A), the cross-talk between two negative feedback loops can be modeled using logical AND or OR operations. (C) The kinetic details that may affect DoRA, depending on modeling choices that are explored in this work. (D) Quantitative metrics for DoRA: The upper left panel shows dose-response curves of the fractional activation for receptors (red) and downstream GTPases (blue: mGTPase; green: tGTPase); these curves after normalization by their activation levels (defined as the level when the stimulus goes to infinity) are denoted by *y*_*Receptor*_ and *y*_*m*(*or t*)*GTPase*_, as shown in the upper right panel; the lower panel defines the DoRA metric as the weighted sum of the distance between *y*_*Receptor*_ and *y*_*m*(*or t*)*GTPase*_. This metric is equal to the area between the *y*_*m*(*or t*)*GTPase*_ and the diagonal line in the plot *y*_*m*(*or t*)*GTPase*_ vs *y*_*Receptor*_. Smaller distance indicates better DoRA.

## 2 Model development

We briefly describe the biochemical circuit that we study here; this circuit was originally described in mammalian cells and experimentally interrogated in (33), (Figure 1A). In the presence of epidermal growth factor (EGF), the active Ras-superfamily mGTPases Arf1 on Golgi membrane recruit GIV/Girdin (a protein that is known to fuel aggressive traits in diverse cancers), and the latter works as guanine nucleotide exchange factor (GEF) to turn tGTPases Gi on (39–43). Subsequently, GIV increases the level of the GAP for mGTPase by molecular scaffolding action, and the activated tGTPase acts as a co-factor to maximally activate the GAP. This circuit was subsequently modeled for cell secretion (44) and for stability analysis (43).

To translate this circuit into a mathematical model, we make the following assumptions. The total number of active and inactive receptors is assumed to be a constant, and so is the total number of mGTPases and that of tGTPases. Based on these assumptions, we choose the fractional activation, which is defined as the ratio of the number in the active form to the total number, to describe the state of the receptor or GTPase. The fraction of the inactive form is one minus the state variable.

### 2.1 Governing equations

The dynamics of mGTPases, tGTPases, and corresponding GEFs and GAPs can be governed by the following system of equations.

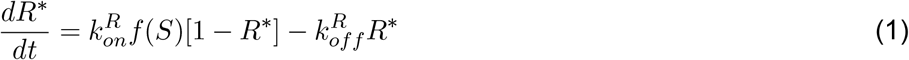

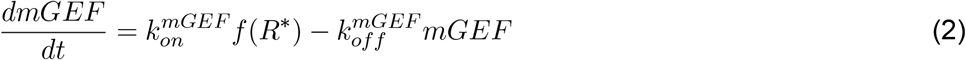

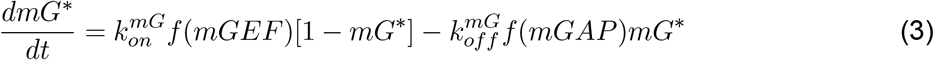

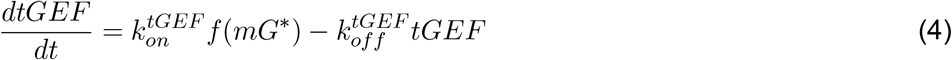

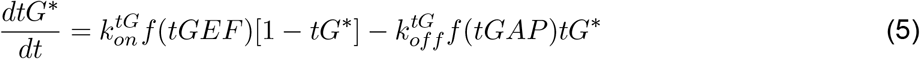

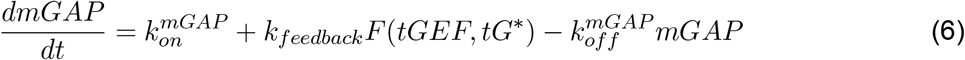

where *S* denotes the stimulus EGF. *R*^*∗*^, *mG*^*∗*^, *tG*^*∗*^ represent the fractional activation of the receptor, mGTPase, and tGTPase, respectively. *mGEF, tGEF, mGAP* are concentrations of GEF for mGTPase (mGEF), GEF for tGTPase (i.e., GIV; denoted as tGEF), GAP for mGTPase (mGAP), respectively. *k*_*on*_’s are production rate constants, *k*_*off*_ ‘s are decay rate constants. The negative feedback loops from the active tGTPase and tGEF to mGAP are modeled by *k*_*feedback*_*F* (*tGEF, tG*^*∗*^), where *k*_*feedback*_ indicates the negative feedback strength and *F* (*tGEF, tG*^*∗*^) the crosstalk between tGEF and active tGTPase. The function *f* describes the effect of the regulator.

#### Form of reaction kinetics *f*

The outcome of a model for signal transduction depends on the form of the reaction kinetics. Here, we explore the coupled GTPase circuit using two classic forms of reaction kinetics – mass action and Hill functions. When the model is developed using mass action kinetics *f* in Eq. (1)-Eq. (6), the *f* becomes

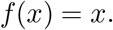

For the Hill-function kinetics,

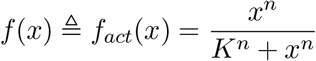

Here *K* and *n* are half-maximal activation and Hill coefficient, respectively.

#### AND and OR logic gates to model feedback term *F* (*tGEF, tG*^*∗*^)

While it is known that tGEF and active tGTPase both exert act mGAP through negative feedback, where they are both required or whether either on is sufficient is not yet clear. Therefore, we consider two options for the feedback by studying the AND logic gate and the OR logic gate. If the AND logic gate is used to model this interaction, *f* (*tGEF*) and *f* (*tG*^*∗*^) are multiplied together; if the OR logic gate is applied, these two terms are added together. As a result, we can write the *F* (*tGEF, tG*^*∗*^) as follows:

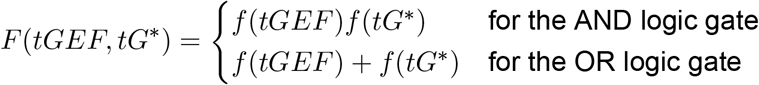

To investigate the role of the negative feedback loops on the DoRA, we also perturbed this circuit by deleting the negative feedback loops (Figure 1B), and this is achieved by setting *F* (*tGEF, tG*^*∗*^) ≡ 0.

Thus, we have six models according to different combinations of *f* and *F* (*tGEF, tG*^*∗*^), because *f* and *F* (*tGEF, tG*^*∗*^) have two and three choices, respectively. The three choices of *F* (*tGEF, tG*^*∗*^), i.e., 0, *f* (*tGEF*)*f* (*tG*^*∗*^) or *f* (*tGEF*) + *f* (*tG*^*∗*^), correspond to the circuit without feedback, the circuit whose negative feedback is modeled by AND logic gate, and the circuit where the OR logic gate is applied. For each circuit, *f* has two choices, indicating which kinetics is adopted. See Supplement section 1 for the numerical simulations for these models.

### 2.2 Metrics for DoRA

The dose-response alignment for species downstream of the receptor can be obtained from inspecting how well dose-response curve of the downstream species is aligned with (or closely matches) the receptor occupancy curve. Quantitatively, this can be measured by the distance between the “normalized” GTPase and the “normalized” receptor dose-response curves (Figure 1D). “Normalized” refers to the fact that the dose-response curve is divided by its maximal fractional activation and ensures that every species falls into the same range [0, 1]. The distance between these two curves is defined as the weighted sum of the absolute deviations (the bottom panel in Figure 1D) and is equal to the area between the diagonal line and the curve of the normalized GTPase response versus the normalized receptor response. If this metric is 0, the normalized fractional activation of GTPase is equal to that of the receptor indicating an exact match of normalized dose-response curves between GTPase and the receptor, i.e., the perfect DoRA is achieved. Since there are two types of GTPase, m- and tGTPase, we calculated this metric for both GTPase (the blue and green curves in Figure 1D). This definition is the integral of the absolute deviations in the vertical direction, while the SWRMS distance defined by Andrews et al. (9) is the integral of the squared deviations in both vertical and horizontal directions. One advantage of the metric used in our work is that it makes detailed theoretical analyses feasible due to its simple form.

## 3 Results

### 3.1 mGTPase activation levels impact DoRA in the absence of negative feedback

We begin our analysis with the simple case of coupled switches without any negative feedback (Figure 1B) for both mass action and Hill-type kinetics. Yu *et al*. showed that when the downstream response has a small value, the system achieves good DoRA (4). Therefore, we first investigated how GTPase activation levels determine GTPases’ DoRA behavior. Here, we only considered the mGTPase activation level, because the positive regulation from mGTPase to tGTPase leads to a positive correlation between m- and tGTPase activation levels.

First, in the mass action model, the DoRA can be improved by decreasing the mGTPase level. The normalized steady-state values of *mG*^*∗*^ and *tG*^*∗*^ are given by (See Supplement section 2.1 for details)

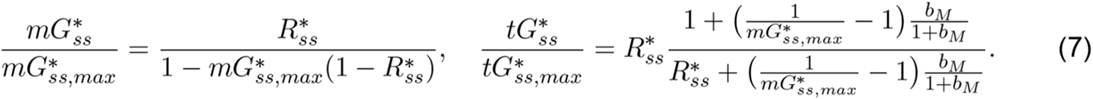

where the subscripts *ss* and *max* denote the steady-state value and the maximal value, respectively. 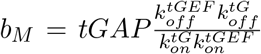, which is independent with the mGTPase activation level 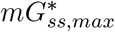 (See also the first plot in Figure 1D). According to these two equations, the normalized m- and tGTPases, i.e., 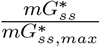 and 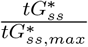, both have two important properties (Figure 2A): the first is that they are always larger than 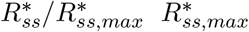 is 1 in the mass action model), and the second is that their values decrease with decreased mGTPase activation level 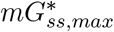. These two properties suggest that the small mGTPase activation level lowers the normalized m- and tGTPase curves and thus makes them close to the receptor curve, resulting in good DoRA behavior for both GTPases. Furthermore, if we tune one kinetic parameter to lower the mGTPase activation level, the DoRA for both GTPases can be enhanced (Figure 2A).

**Figure 2:**
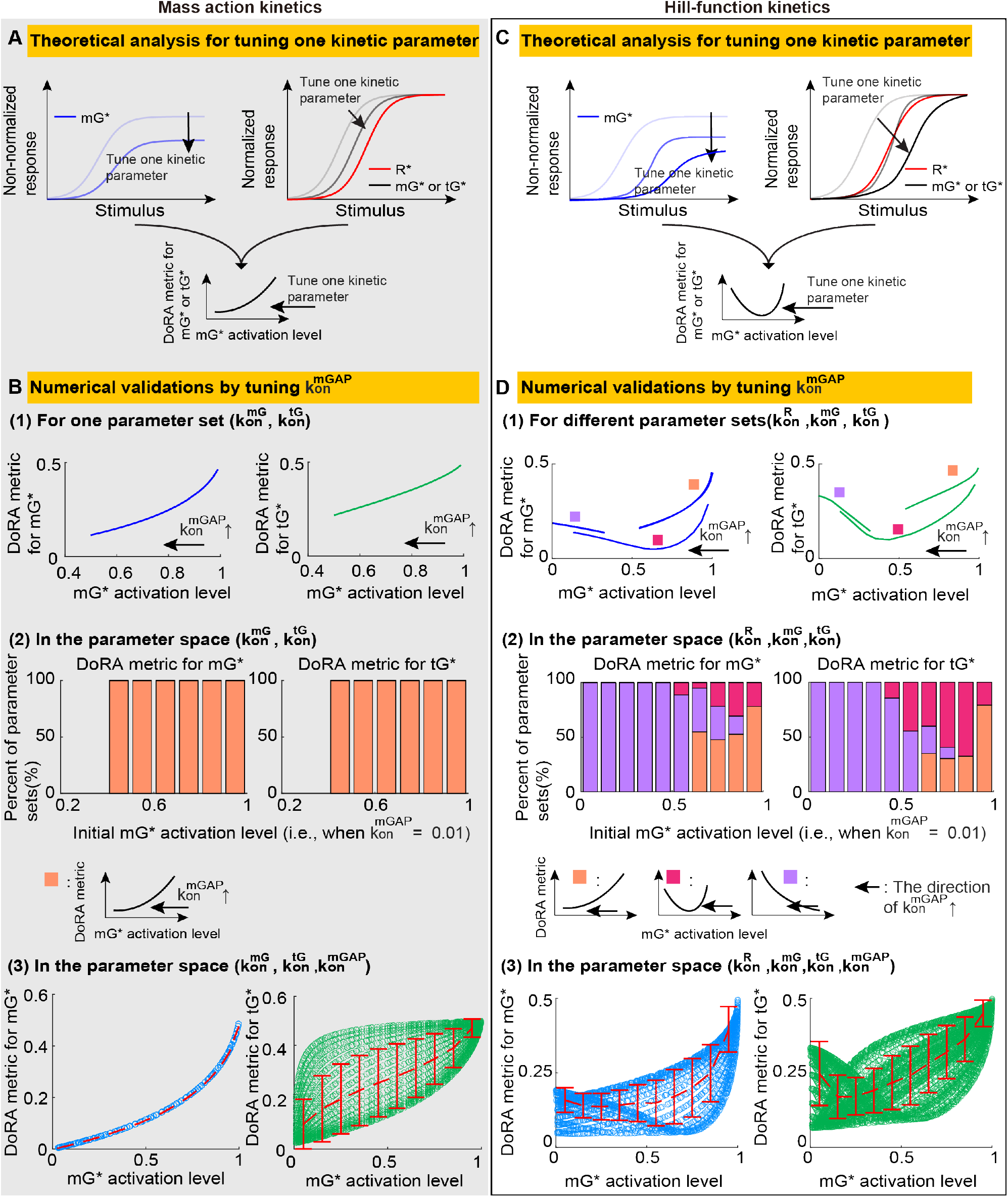
The relations between the DoRA behavior and the mGTPase activation level in the circuit without feedback. (A) The DoRA behavior improves with the decreased mGTPase activation level when tuning one kinetic parameter in the mass action model. (B) Numerical validations for results in A by tuning 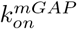. The upper panel: the DoRA metric vs the mGTPase activation level when increasing 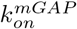 (denoted by 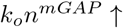) for a given parameter set. The middle panel: percents of maintaining the trend in the upper panel when searching the parameter space 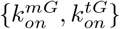. The bottom panel: the DoRA metric vs the mGTPase activation level in the parameter space 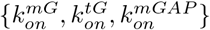, with the dashed red line the mean and the error bar the standard deviation. See Table S1 for parameters. (C) The DoRA behavior improves and then becomes bad with the decreased mGTPase activation level when tuning one kinetic parameter in the Hill-function model. (D) Same plot as in B except that the Hill-functions kinetics is adopted. The parameter space also expands to include the 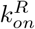, because the 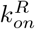 affects the receptor activation level and thus the mGTPase activation level in the Hill-function model.

We further validated the above analysis numerically by studying the effects of only tuning 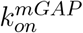. For a given parameter set, increasing 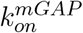 leads to not only the decrease in mGTPase activation level but also the improved DoRA behavior for both GTPases (the upper panel in Figure 2B; also see Figure S1A for dose-response curves). This trend is not affected by the choice of parameter sets: for every 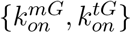 set, which may have different mGTPase activation levels when the 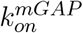 is 0.01, increasing 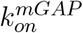 from 0.01 to 1 causes the decrease of both the DoRA metric and the mGTPase activation level (the middle panel in Figure 2B). The above results indicate that when other kinetic parameters are fixed, increasing 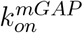, i.e., decreasing mGTPase activation level, can enhance DoRA for both GTPases. This conclusion is not influenced by choosing which kinetic parameter to tune (Figure S2). Furthermore, to study the effects of tuning more than two kinetic parameters simultaneously instead of only tuning 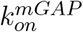, we plotted the scatter plots of the mGTPase activation level vs the DoRA metric in the parameter space 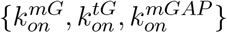 (the bottom panel in Figure 2B). The DoRA behavior of mGTPase can be improved as long as the mGTPase activation level is decreased, because there exists a one-to-one mapping from the mGTPase activation level to the mGTPase’s DoRA metric. Nevertheless, decreasing the mGTPase activation level can impair or improve the tGTPase’s DoRA behavior due to the one-to-many mapping, but the improvement effect has a larger probability to occur according to the mean DoRA trend.

Next, we turned to the Hill-function model and found that an intermediate mGTPase activation level benefits the DoRA for the circuit without feedback. In the Hill-function model, the mGTPase activation level, i.e., the maximal value of the active mGTPase under varied stimuli, is given by:

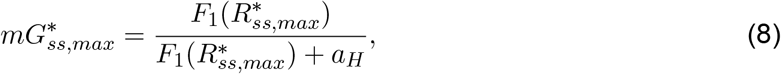

and the normalized steady-state values of *mG*^*∗*^ and *tG*^*∗*^ are shown as follows (See Supplement section 2.2 for details)

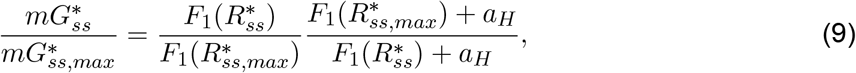

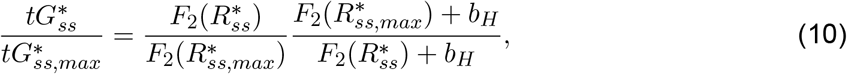

where 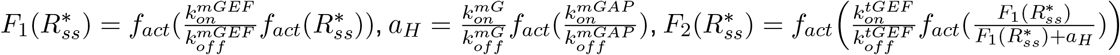 and 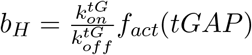. The subscripts *ss* and *max* have the same definitions as before. Unlike the case for the mass action model, it is hard to rewrite 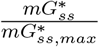 as a function of 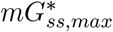. This prevents us from directly obtain how 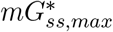 affects 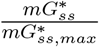. However, we can vary one kinetic parameter and study the corresponding changes of 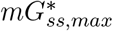 and 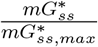. The varied kinetic parameter can be 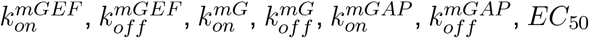, or *B*, but we didn’t consider the last two kinetic parameters due to their complexity (*EC*_50_ and *B* can be different for distinct *f*_*act*_). By deriving derivatives of 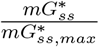 and 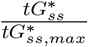 with respect to each kinetic parameter (Eq. (S12)), we proved that 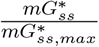 and 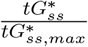 will decrease when one kinetic parameter is tuned to reduce the 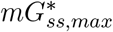. In other words, the normalized m- and tGTPase curves are decreased if one kinetic parameter is changed to decrease the mGTPase activation level (Figure 2C). However, the normalized GTPases curves are not always larger than that of the receptor due to the nonlinearity of the *F*_1_, and thus the normalized GTPase curves may cross the receptor curve from the left to the right, suggesting that the DoRA behavior improves and then worse as the mGTPase activation level decreases (Figure 2C). Therefore, an intermediate mGTPase activation level results in good DoRA for the Hill-function modeled circuit in the absence of feedback.

Similar to the numerical validations in the mass action model, we also took 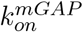 as an example to verify the above analysis for the Hill-function model. For three different parameter sets of 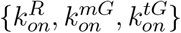, increasing the 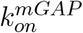 from 0.01 to 1 all reduces the mGTPase activation level, but the DoRA metric for both GTPases can be decreasing, increasing, or decreasing at first and then increasing (the upper panel in Figure 2D; also see Figure S1B). Note that the decreasing trend of the DoRA metric with increased 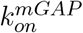 tends to have large mGTPase activation levels, the increasing trend the small mGTPase activation levels, and the non-monotonic trend the intermediate mGTPase activation levels. This also holds when we sampled more parameter sets of 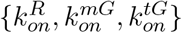 in the whole parameter space (the middle panel in Figure 2D): the decreasing, increasing, and non-monotonic trends are located in high, low, and intermediate initial mGTPase activation level (the value when 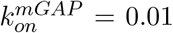, respectively. These simulations indicate that an intermediate mGTPase activation is preferred for the DoRA behavior, which also holds when tuning other kinetic parameter rather than 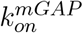 (Figure S3). Furthermore, in the parameter space 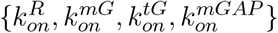, the relation between the mean DoRA behavior and the mGTPase activation level shows consistent results (the bottom panel in Figure 2D).

### 3.2 The effect of negative feedback is model-dependent and mGTPase activation level-dependent

The above analyses focused on the coupled switches without feedback. Next, we will investigate the effect of adding negative feedback. Although the effects of the feedback is the same as the high 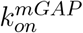 value for the circuit without feedback based on their inhibitory roles in the mGTPase activation level, their effects on the DoRA may differ a lot because the feedback induces more non-linearity. The strength of the feedback is tuned by varying *k*_*feedback*_, where 0 means no feedback and a large value indicates strong feedback. Moreover, as we have shown in the previous section, different kinetics forms lead to different DoRA performance, so the mass action and Hill-function kinetics are both considered.

In the mass action model, adding negative feedback has diverse effects on DoRA: the negative feedback enhances (or impairs) DoRA when the mGTPase activation level is high (or low), while the intermediate mGTPase activation level leads to the non-monotonic effect of negative feedback. The theoretical analyses for the AND logic gate are taken as an example and summarized as follows (See Supplement section 3.1 for details). The steady state of the fractional activation of mGTPase satisfies the following equation:

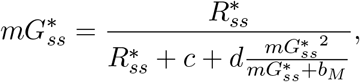

where *d, c* and *b*_*M*_ are constants, and the *k*_*feedback*_ is only involved in *d*. According to this equation, the 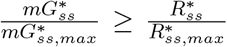 all the time, suggesting a higher location of normalized mGTPase dose-response curve than the receptor curve. Besides, the derivative of 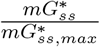 with respect to the *k*_*feedback*_ is shown as follows (See Supplement):

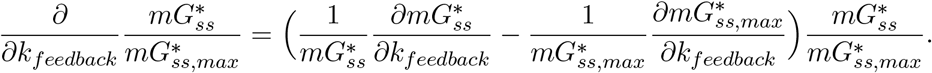

The term in the bracket is the difference of the function 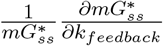 at 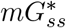 and 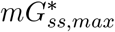. Since this function decreases at first and then increases with increased 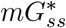 (Figures S4-S7, and Eq. (S13)), the sign of 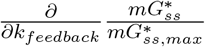 is positive for small 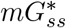 and negative for large 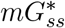. In other words, the small (or large) 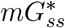 leads to the increase (or decrease) of the normalized curve when increasing feedback strength (gray curves in 3A). Therefore, if the mGTPase activation level is high enough, the decreasing trend of the normalized mGTPase curve with increased feedback strength dominates, enhancing the DoRA for mGTPase (➀ in Figure 3A; also see Figure S4). Similarly, when the mGTPase activation level is low, the strong feedback impairs the DoRA for mGTPase (➁ in Figure 3A; also see Figure S5). When the mGTPase activation level is neither high nor low, non-monotonic effect occurs. Similar conclusions can be drawn for tGTPase (See Supplement).

**Figure 3:**
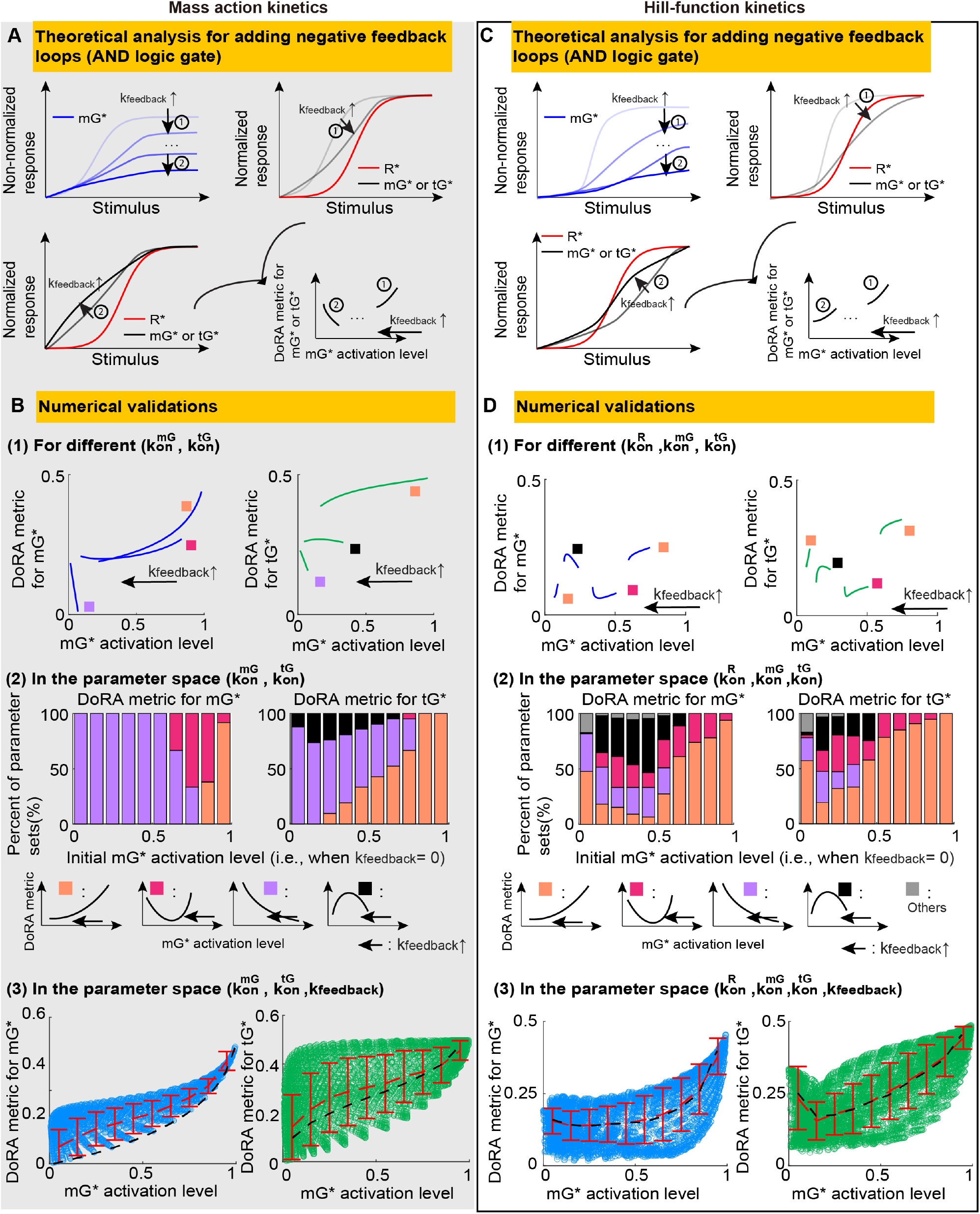
Effects of adding the negative feedback. Same plot as in Figure 2, except that the feedback strength *k*_*feedback*_ is tuned while maintaining other kinetic parameters unchanged. The three dots in A indicate the region where the non-monotonic change of the DoRA metric may occur. The black dashed lines in the lower panels in B and D are the red dashed lines in the lower panels of Figure 1B and 1D, respectively. Here, the AND logic gate is used to model the negative feedback loops. See Table S1 for parameters.

We also validated above conclusions through numerical simulations. For different values of 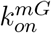 and 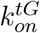, increasing feedback strength (i.e., increasing *k*_*feedbak*_ from 0 to 10^2.6^) all causes the decrease of the mGTPase activation level, but trends for the DoRA metric are diverse (upper and middle panels in Figure 3B; also see Figure S1C): increasing (purple), decreasing (orange), decreasing first and then increasing (magenta), or increasing first and then decreasing (black). Furthermore, increasing and decreasing trends usually occur when the mGTPase activation level in the absence of feedback is low and high, respectively. These results are consistent with the theoretical analyses. Though we elaborated on AND logic gate, the results are the same when using OR logic gate (Figure S8A-B).

Another property of adding the negative feedback loops in the mass action model is that it causes a big change in the DoRA metric (the bottom panel in Figure 3B). In this plot, red and black dashed lines denote the mean DoRA metric for the circuit with and without feedback in the parameter space respectively, where the black dashed lines are the red lines in the bottom panel in Figure 2B. We can find that, the mean DoRA metric for both GTPase change a lot after adding the negative feedback, indicating a large effect of the negative feedback on DoRA in the mass action model. This is also supported by the large change of DoRA metric in the upper panel in Figure 3B. Moreover, we note that the mean DoRA metric in the presence of negative feedback is higher than that without feedback, i.e., the circuit with negative feedback shows worse DoRA performance than the circuit without feedback under the constraint of the same mGTPase activation level. This result does not violate the widely accepted principle that negative feedback improves DoRA. Adding or strengthening the negative feedback for a given parameter set reduces the mGTPase activation level and thus decreases the mean DoRA metric, resulting in a high probability of generating good DoRA.

Next, we turned to the Hill-function model to explore the role of negative feedback. Unlike the case in the mass action model, adding negative feedback in the Hill-function model usually enhances the DoRA behavior when the mGTPase activation level is high or low. The steady state of the fractional fractional of the mGTPase for AND logic gate satisfies the following equation (See Supplement section 3.2 for details):

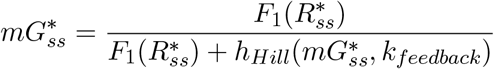

where 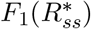 has been defined in the previous section. The 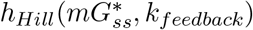 represents the contributions of tGTPases and tGEF towards the mGAP. This equation cannot ensure that the normalized curve for mGTPase is higher than that for receptor due to the non-linearity of the function *F*_1_. In fact, the high mGTPase activation level usually corresponds to a higher location of the mGTPase curve than the receptor curve, and increasing *k*_*feedback*_ under this condition decreases the location of the mGTPase curve (See Supplement section 3.2), leading to a good DoRA of mGTPase (the ➀ in Figure 3C). On the other hand, the low mGTPase activation level often means a lower location of the mGTPase curve than the receptor curve, and increasing *k*_*feedback*_ at this time raises the mGTPase curve (See Supplement section 3.2), also ensuring the positive role of negative feedback for the DoRA of mGTPase (the ➁; in Figure 3C). Similar conclusions can be drawn for tGTPase (See Supplement section 3.2).

Numerical simulations support the above conclusions. Here, we demonstrated the AND logic gate to model the negative feedback loops, while the OR logic gate shows similar results (Figure S8C-E). For two given parameter sets with either high or low mGTPase activation level (marked by orange boxes in the upper panel in Figure 3D; Figure S1D), increasing the feedback strength, i.e., varying *k*_*feedback*_ from 0 to 10^2.6^, decreases the DoRA metric; in the whole parameter space 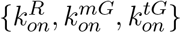, the high or extremely low mGTPase activation level often leads to the decreasing trend of the DoRA metric (orange bars in the middle panel in Figure 3D). Since the mass action kinetics shows such decreasing trend of the DoRA metric only when the mGTPase activation level is high, these results suggest that the Hill-function kinetics has a larger parameter space to produce this decreasing trend. Therefore, compared with the mass action model, the effect of negative feedback in the Hill-function model is more unified improving DoRA in most cases. However, when the mGTPase activation level is in the intermediate range, trends of the DoRA metric with respect to the feedback strength are non-monotonic and diverse (magenta, black and gray colors in Figure 3D; Figure S1D).

The overall effects of negative feedback in the Hill-function model are shown in the lower panel in Figure 3D. The red and black dashed lines still represent the mean DoRA metric with and without negative feedback in the parameter space 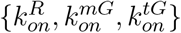, respectively. Unlike the case in the mass action model where two lines are far from each other, these two curves are almost overlapped in the Hill-function model, indicating a small effect of negative feedback on the DoRA. This may result from the saturation effect of the Hill function; the whole system cannot respond to further changes in the feedback strength when the system is saturated.

### 3.3 The OR logic gate has the similar DoRA behavior to the AND logic gate

While modeling the negative feedback loops with the AND or OR logic gate does not change the trend of the DoRA with increased feedback strength, the DoRA behavior with a certain feedback strength may differ. To make a fair comparison, for the system with the AND or OR logic gate, we chose the same parameter set but allowed different values of *k*_*feedback*_ to ensure the same *mG*^*∗*^ activation levels. This constraint is based on the importance of the mGTPase activation level as shown in previous sections. Once we have such kinetic parameters and the different *k*_*feedback*_, we computed and compared the DoRA metrics between these two systems (corresponding to one dot in Figure 4A). After we randomly chose kinetic parameter sets and corresponding *k*_*feedback*_, we can compare several pairs of DoRA metrics for the systems using different logic gates (Figure 4A).

**Figure 4:**
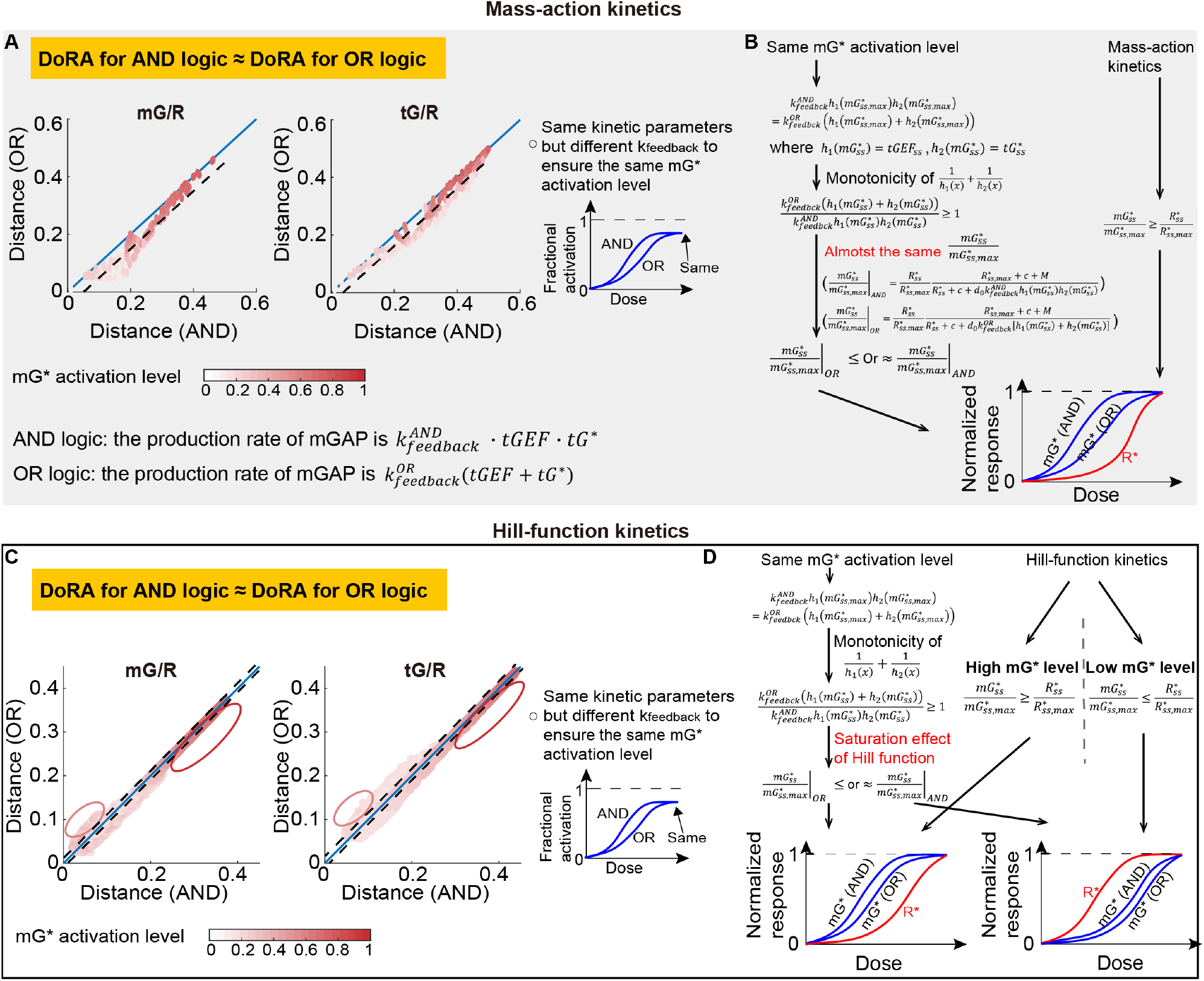
The OR logic gate has the similar DoRA behavior to the AND logic gate. (A) Comparisons of the DoRA metrics between the system with the AND logic gate and that with the OR logic gate. The AND logic gate means that the production rate of mGAP is modeled by 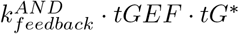 ; the OR logic gate corresponds to 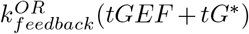. Each dot corresponds to one kinetic parameter set in the parameter space 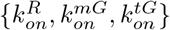, and the 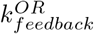 is different from 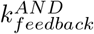 to ensure the same *mG*^*∗*^ activation levels (the value when the stimulus *S*→ ℞). The black dashed line indicates the averaged distance from dots to the diagonal line. The diagonal line is in blue. (B) Theoretical analysis for comparisons of DoRA metrics between AND and OR logic gates in the mass action model. The *tGEF* can be rewritten as a function of *mG*^*∗*^, i.e., *h*_1_(*mG*^*∗*^); the *tG*^*∗*^ is rewritten as the function *h*_2_(*mG*^*∗*^). (C-D) The same plots as in A-B but the Hill-function kinetics is adopted.

Interestingly, in the mass action model, the AND and OR logic gates show almost the same DoRA behavior. This is demonstrated by the small distance between the diagonal line and dots in Figure 4A, where x and y coordinates of these dots are the DoRA metric for the AND and OR logic gates, respectively. Next, we validated this finding theoretically. Though the OR logic gate shows better DoRA performance than AND logic gate (See Supplement section 4 and Eq. (S14)), this advantage of OR logic gate is negligible if the feedback strength is close to zero (the dots in dark red in Figure 4A), because the model using the OR or AND logic gate both degenerates to the same model in the absence of the feedback. When the feedback strength is nonzero, it seems that the DoRA performance for the AND logic gate and that for the OR logic gate are still almost the same. This might come from the almost same 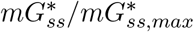 for both logic gates (Figure 4B):

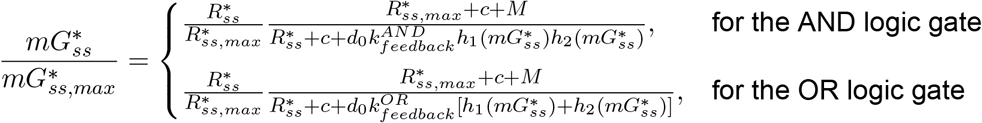

Where 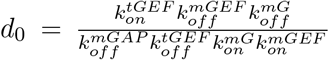, and *c* has the same definition as in previous sections. The 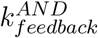 and 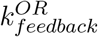 represent the negative feedback strength for the system with the AND logic gate and the system with the OR logic gate, respectively. The 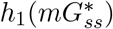 and 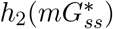 describe how *mG*_*ss*_ determines *tGEF*_*ss*_ and 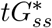, respectively. 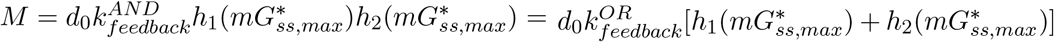, where the last equality originates from the constraint of the same mGTPase activation level. The only difference in the last term in the denominator cannot have a huge impact on the 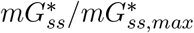 value.

As for the Hill-function model, the DoRA behavior for the OR logic gate is still almost the same as that for the AND logic gate (Figure 4C). The fact that the 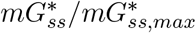 curve for the AND logic gate is higher than that for the OR logic gate still holds (the left column in Figure 4D), but the 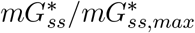 curve moves from the left to the right of the receptor curve with increased feedback strength in the Hill-function model (the right column in Figure 4D). Therefore, the high mGTPase activation level (i.e., low feedback strength) results in better DoRA for the OR logic gate than the AND logic gate (dots in the red box in Figure 4C); the low mGTPase activation level may lead to the better DoRA for the AND logic gate (dots in the pink box in Figure 4C). But one important feature of these dots is that they are more close to the diagonal line, compared with those in the mass action model. This is because of the saturation effect of Hill function: though the mGAP levels in the model using AND or OR logic gate can differ a lot, its effect on the mGTPase, *f*_*act*_(*mGAP*), might be the same if *f*_*act*_ reaches the saturation level 1.

### 3.4 DoRA in a longer cascade can be obtained using Hill-function kinetics under certain conditions

From above analyses, the DoRA behaviors for m- and tGTPases always show the same tendency, for example, when increasing mGAP level, adding negative feedback, or modeling feedback with distinct logic gates. But which GTPases have a better DoRA performance under the same circumstance? To answer this question, we compared the DoRA metrics between m- and tGTPases in the mass action or Hill-function model.

For the mass action model, the DoRA behavior for tGTPase is worse than that for mGTPase, suggesting that longer cascade shows worse DoRA performance. This result is validated by the following theoretical analyses (Figure 5A). As we have shown in the previous section and the supplement, the normalized steady-state value of *mG*^*∗*^ in the circuit without feedback is given by

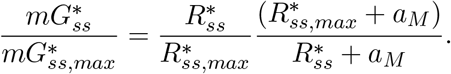

**Figure 5:**
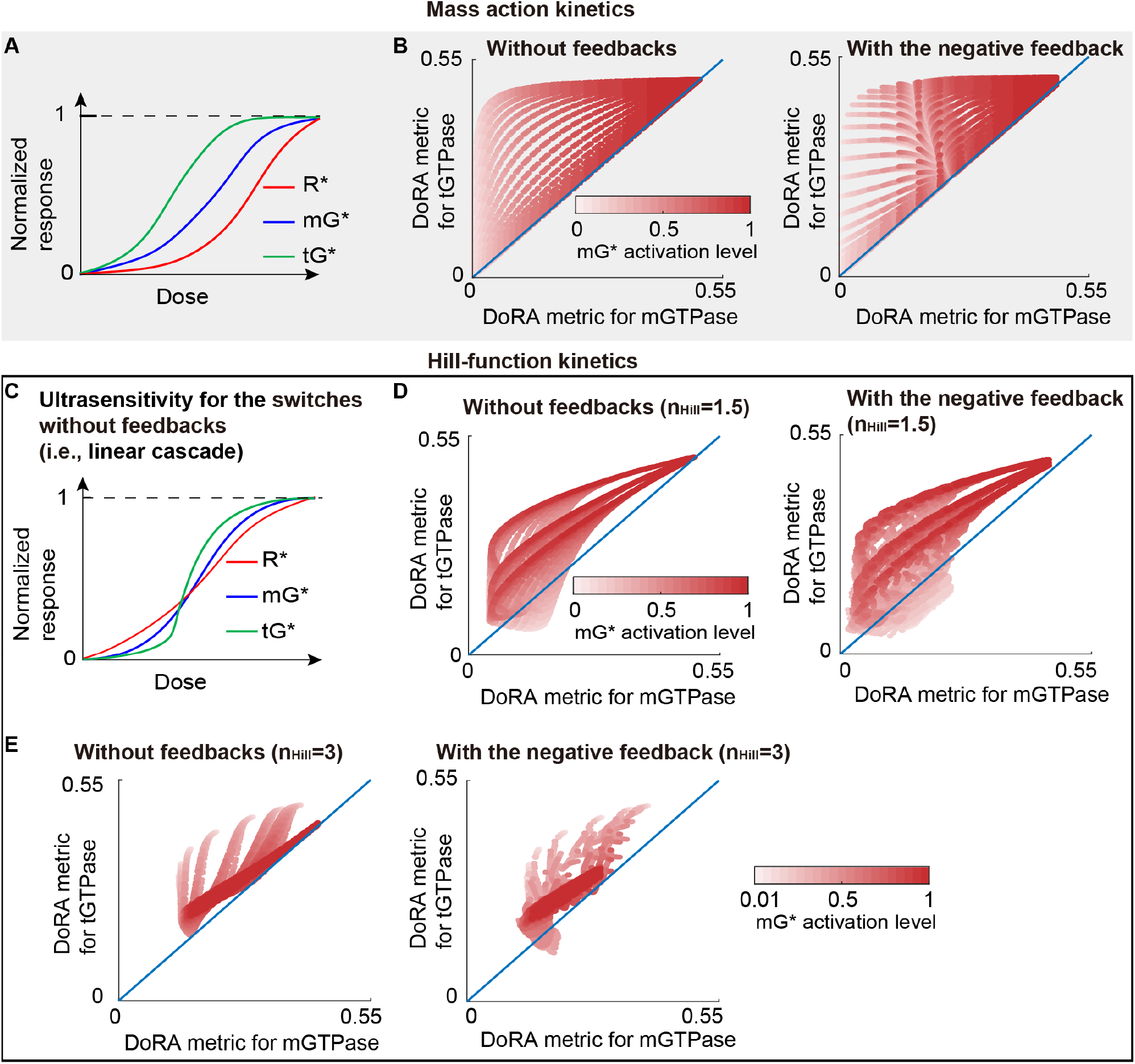
DoRA in a longer cascade can be obtained using Hill-function kinetics and low mGTPase activation level. (A) The DoRA of mGTPase is better than that of tGTPase in the mass action model. This is because 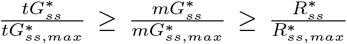 holds all the time. (B) The scatter plots of DoRA metric for mGTPase vs tGTPase in the mass action model. Each dot corresponds to a set of kinetic parameters. The blue line is the diagonal line. The left and right panels correspond to the case without and without feedback, respectively. (C) The ultrasensitivity for the circuit without feedback where the Hill-function kinetics is adopted. (D-E) Same plots as in B except that the model is Hill-function based with the Hill coefficient 1.5 (D) or 3 (E).

Thus, 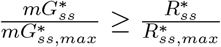 all the time due to 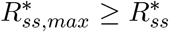. For the circuit with negative feedback loops, *a*_*M*_ becomes an increasing function of 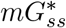, and thus this relation also holds. Using a similar way, we can prove that the 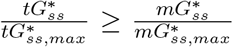 all the time. Therefore, we got 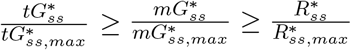, that is, the normalized dose-response curve of tGTPase is higher than that of mGTPase, and the latter is higher than that of the receptor. According to the locations of different curves, the tGTPase curve is farther from the receptor curve, leading to the worse DoRA. This is also supported by the scatter plots of the DoRA metric between mGTPase and tGTPase (Figure 5B).

As for the Hill-function model, the better DoRA performance of tGTPase than mGTPase occurs when the mGTPase activation level is low. A widely observed property in the linear cascade is the ultrasensitivity, where the long cascade results in a steep response curve (Figure 5C). It means that the mGTPase curve is closer to the receptor curve than the tGTPase curve, especially when the dose is very low or high. Thus, the mGTPase exhibits better DoRA behavior than the tGTPase in most cases (dots above the diagonal line in Figure 5D and S8F). Other cases where the tGTPase performs better may result from the complicated behavior when the dose is in the intermediate level; numerical simulations suggest that these cases may occur when the mGTPase activation level is low (dots under the diagonal line in Figure 5D and S8F). The underlying mechanism for why tGTPase can display a better DoRA behavior with a low mGTPase activation level remains unknown, and increasing the Hill coefficient to 3 does not seem to expand this advantage. Moreover, increasing the Hill coefficient makes the lowest value of DoRA metrics increase (Figure 5D-E and Figure S9), indicating that a high Hill coefficient may impair DoRA.

## 4 Discussion

A longstanding question in biology is how cells accurately transmit information from the extracellular space and thus make right decisions. The DoRA is one of the mechanisms to ensure cells respond proportionally to the external stimulus. Though the “push-pull” topology can produce perfect DoRA (7, 9), the prevalence of the negative within DoRA motifs may indicate the possibility of the negative feedback to achieve the near-perfect DoRA under certain circumstances. We explored the design principles of DoRA in the coupled GTPase switches, where the negative feedback loops couple two types of GTPase switches. By varying the kinetic form, the concentration of mGAP, the strength of the feedback, the logic gate for two feedback loops, the length of the cascade, or the Hill coefficient, we obtained the following (summarized in Figure 6):

- For coupled switches without feedback, kinetic parameters that ensure low mGTPase activation produce good DoRA in the mass action model (black solid line in Figure 6); however, an intermediate mGTPase activation level is preferred in the Hill-function model (red solid line in Figure 6).
- Adding negative feedback loops cannot always enhance the DoRA; the enhancement role of negative feedback loops occurs for the mass action model with high mGTPase activation (the right black arrow in Figure 6), or for the Hill-function model with high or extremely low mGTPase activation, i.e., a broader parameter space than that in the mass actions model (red arrows in Figure 6).
- Adding negative feedback loops causes a larger change in DoRA performance in the mass action model compared with the Hill-function model.
- Modeling two negative feedback loops *tGEF* → *mGAP* and *tG*^*∗*^ → *mGAP* with the AND or OR logic gate shows similar DoRA behavior under the constraint of the same mGTPase activation level (the curved arrow in Figure 6).
- DoRA in a longer cascade can be obtained using Hill-function kinetics and with low mGTPase activation level (dashed arrow in Figure 6).

**Figure 6:**
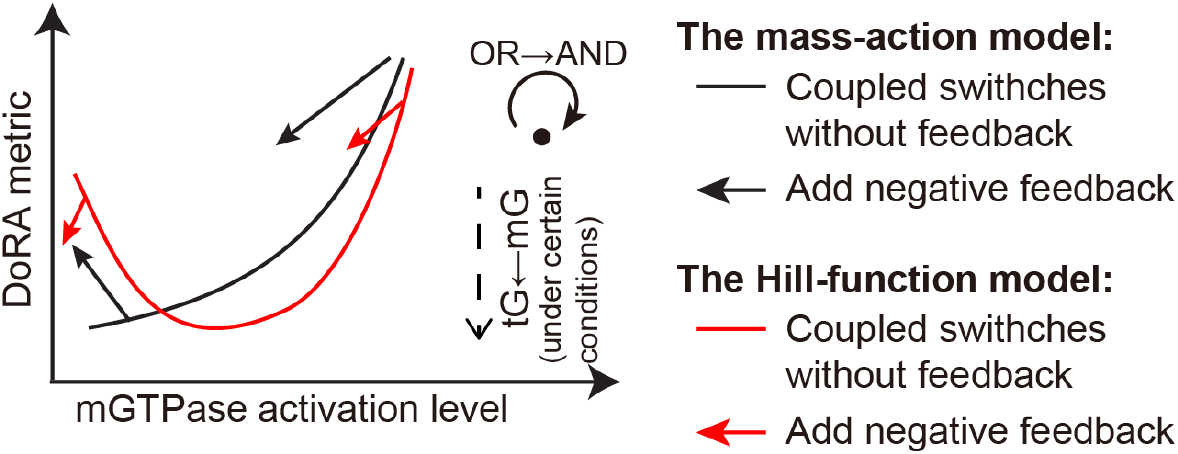
Summary of the effects of kinetic details on the DoRA.

These results suggest the emergence of different DoRA behavior when comparing the simpler mass action kinetics to the more complex (i.e., nonlinear activation rates) Hill-function kinetics. This is mainly because the Hill-function model allows more relative positions of normalized dose-response curves for GTPase and the receptor (Figure 2C and Figure 3C), while the mass action model only exhibits one case–the normalized dose-response curve for GTPase is higher than that for the receptor (Figures 2A and 3A; Fig. 2A in (7) if scaling the curve to ensure the maximal value 1). It appears that the Hill-function kinetics reveal more features than the mass action kinetics. First, the Hill-function model seems to attain DoRA more easily than the mass action model, because the biologically plausible level of the mGTPase activation cannot be extremely low, deviating the condition for good DoRA in the mass action model. Second, the Hill-function model shows a loosened requirement on mGTPase activation level to achieve the negative feedback’s role in improving DoRA than the mass action model; the experiment also observed the negative feedback’s role in DoRA (4).

Through the analysis of three coupled molecular switches (i.e., a longer cascade), we found that the reactant in the distal switch exhibits better DoRA than that in the proximal switch only when Hill-function kinetics is adopted and the mGTPase activation level is low. This suggests another advantage of the Hill function kinetics—making the downstream reactant achieve better DoRA than the upstream reactant. This advantage ensures the information transfer in long cellular signaling pathways. Although the negative feedback loops are widely regarded to linearize the downstream reactant (4, 45), that we observed similar results for the circuit without or with feedback (the bottom panel in Figure 3D) implies that the key factor to maintain a better DoRA for tGTPase might be the level of mGTPase activation instead of the negative feedback, providing new insight about the mechanism for good DoRA.

Finally, this work analyzes DoRA behavior in coupled molecular switches (i.e., GTPases), in which each species is limited to just 2-states (GDP or GTP-bound). One notable distinction between DoRA in coupled switches versus other instances where phosphorylation or other such cascades are in play, is that the 2-states are directly dependent on the local concentrations of GEFs (rate limiting) and GAPs and hence, they are unlikely to be saturated in certain circumstances (e.g. saturation of enzymes in a covalent modification cycle tends to reduce information transmission). By including the GAPs and GEFs in the circuit (46), this work ensures that the emergent features of the motif are explored while attempting to preserve the biological context.

One limitation of our studies is that we neglected the effect of basal constant production rates. The non-zero basal production rates will cause the non-zero response even under the extremely low stimulus, leading to a change of relative positions of dose-response curves. While the basal constant production rates are usually small, we anticipate that our main results will not change dramatically. Another future direction is to study the effect of the receptor number on the DoRA in the circuits we studied. In many signaling pathways (27, 47–51), different receptor abundance cannot affect the downstream response, i.e., the system is robust to the change of the receptor abundance. Therefore, analyzing the mechanism of DoRA achievement under distinct receptor numbers for the circuit with the negative feedback will be significant.

## Acknowledgments

This work was supported by the National Institute of Health Grants: CA238042, AI141630 and CA160911 (to P.G.), GM132106 (to P.R). P.R was also supported by the Air Force Office of Scientific Research (AFOSR) Multidisciplinary University Research Initiative (MURI) Grant FA9550-18-1-0051.

## Supplement

## 1 Numerical simulations

Numerical simulations were implemented in MATLAB. To obtain the dose-response curve for a given parameter set, we use the solver ode15s to simulate the dynamics of Eq. (1)-Eq. (6) on the time interval [0,10^8^] for the stimulus varied from 10^*−*5^ to 10^3^. The large time interval ensures that the steady state is reached. Matlab codes can be accessed at https://github.com/RangamaniLabUCSD/Qiao_et_al_Dose-response-alignment.

## 2 Analysis for the relation between the mGTPase activation level and the DoRA in coupled switches without feedback

### 2.1 The mass action model

We first focused on the coupled switches without feedback that is modeled by the mass action kinetics. In this case, *F* (*tGEF, tG*^*∗*^) = 0 and *f* (*x*) = *x*. Based on these two equations, we set the right terms of Eq. (1)-Eq. (6) equal to zeros and got the following equations:

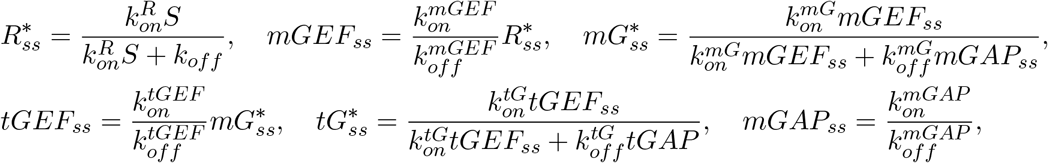

where 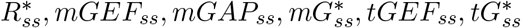, denote the steady-state values of *R*^*∗*^, *mGEF, mGAP, mG*^*∗*^, *tGEF, tG*^*∗*^, respectively, From these equations, we can easily get the following expressions for 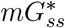 and 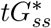

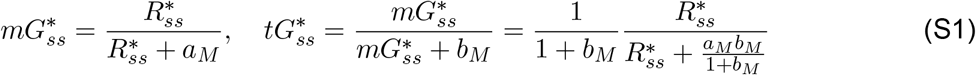

where 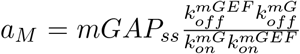 and 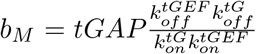. The maximal value of 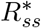 is 1, which is obtained when the stimulus *S* goes to infinity, so the maximal values of 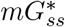 and 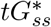 are given by:

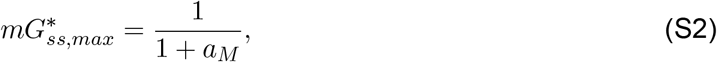

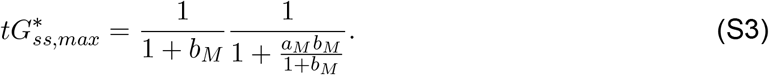

Therefore, the normalized 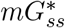 and 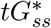 are shown as follows:

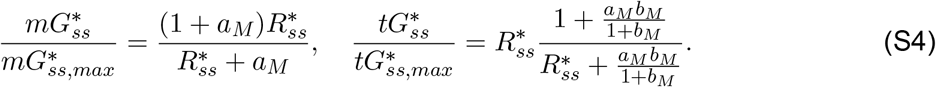

Then, if we rewrite *a*_*M*_ as a function of 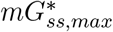 using Eq. (S2) and substitute the *a*_*M*_ in Eq. (S4), we get the following relation between normalized GTPases and 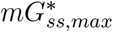:

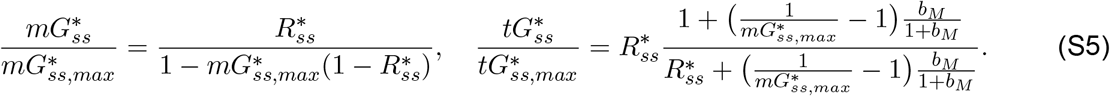

From Eq. (S4) and Eq. (S5), we can obtain following properties for m- and tGTPase:

- Normalized GTPase curves are always higher than the receptor curve, because Eq. (S4) leads to 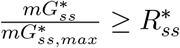 and 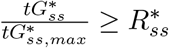;
- Decreasing the mGTPase activation level lowers the normalized GTPase curves, because Eq. (S5) indicates that 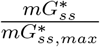 and 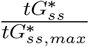 both decrease with decreased 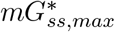.

Taking these two properties together, we conclude that the low mGTPase activation level makes the normalized GTPase curves close to the receptor curve, i.e., good DoRA.

### 2.2 The Hill-function model

Next, for the same circuit, we turned to the Hill-function model to explore the relation between the mGTPase activation level and the DoRA behavior. Here, *f* (*x*) is Hill function, and *F* (*tGEF, tG*^*∗*^) = 0. Let the right-hand terms of Eq. (1)-Eq. (6) equal to zeros and substitute *f* with the Hill function *f*_*act*_. Then relations between steady states are as follows:

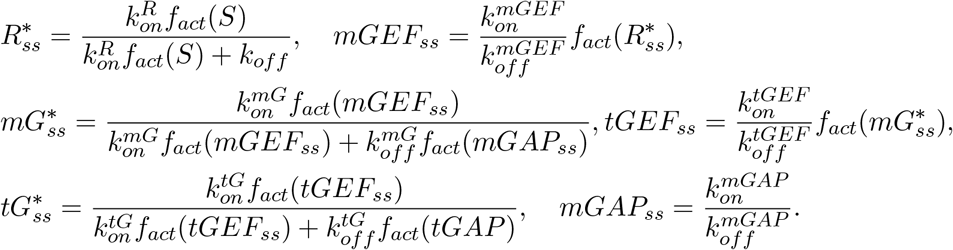

Based on these equations, we got the following equations for steady states of mG* and tG*:

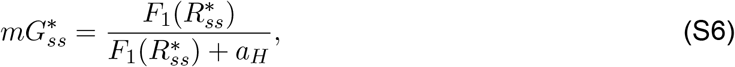

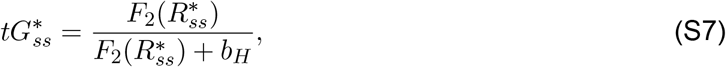

where 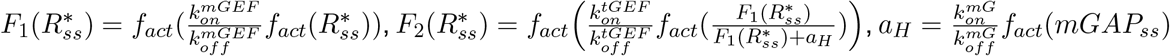 and 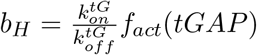. We used 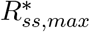 to denote the value of 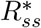 when the stimulus *S* goes to infinity. It should be noted that 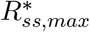 is no longer 1 because the *f*_*act*_(*S*) has the maximal value 1 and thus cannot become infinity even when *S* goes to infinity. So, the maximal values of 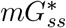 and 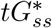, denoted by 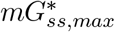 and 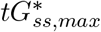, are given by:

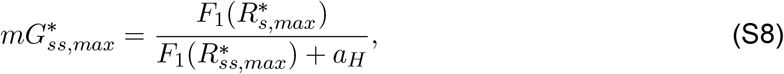

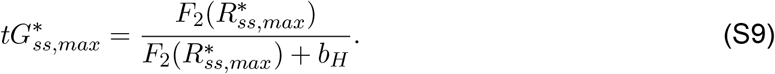

Dividing the 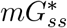 and 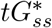 by these two maximal values, we got the following equations:

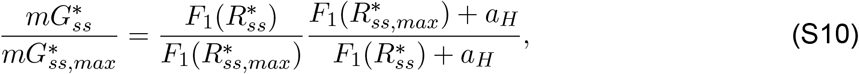

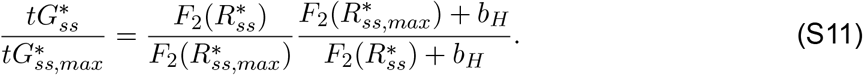

For the DoRA of the mGTPase, tuning one kinetic parameter while fixing other kinetic parameters to reduce the mGTPase level usually leads to the decrease of 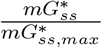. The proof for this conclusion depends on which kinetic parameter is chosen to tune. If 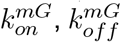, or *mGAP*_*ss*_ is chosen, every term in Eq. (S6) and Eq. (S10) is unchanged except *a*_*H*_; therefore, increasing *a*_*H*_, i.e., decreasing mGTPase activation level 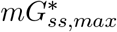, causes the decrease of 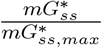. On the other hand, if 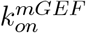, or 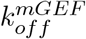 is chosen to tune, the function *F*_1_ in Eq. (S6) and Eq. (S10) changes. The way to tune 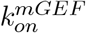 (or 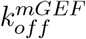) in order to lower the mGTPase activation level is decreasing 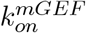 (or increasing 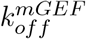), leading to the decrease of 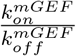. For simplicity, we used *k* to denote 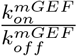. Then we proved that the decrease of *k* results in the decrease of the 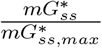 as follows. First, from Eq. (S10) the derivative of 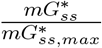 with respect to the *k* (i.e., 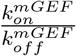) is given by:

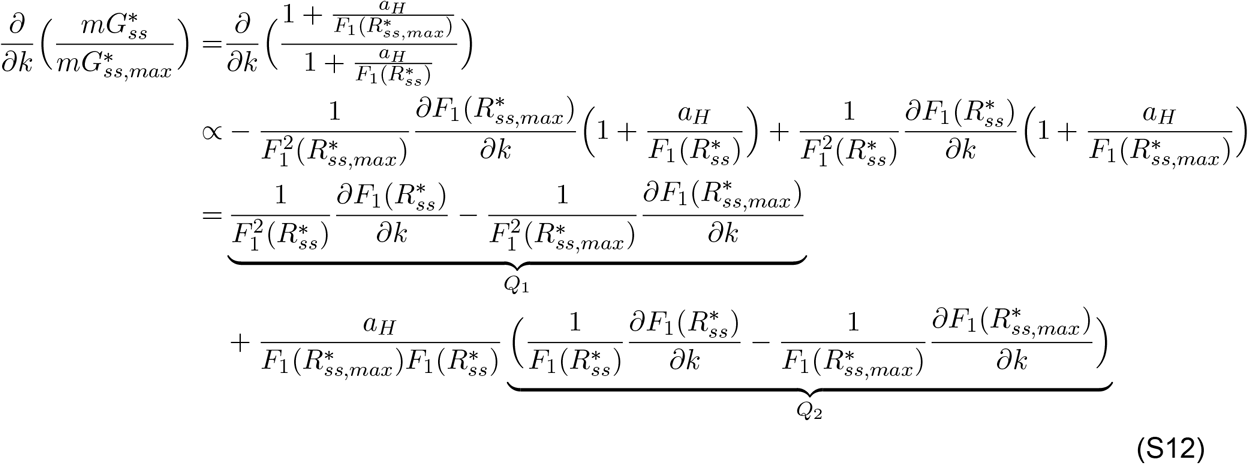

Then, according to the definition of *f*_*act*_, the 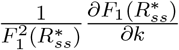 and 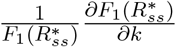 can be rewritten as follows:

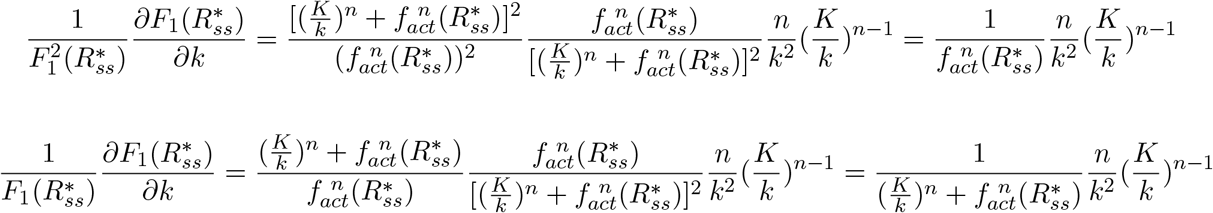

Thus, 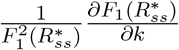 and 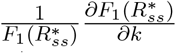 are both decreasing function of 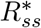, and thus *Q_1_ >* 0 and *Q_2_ >* 0. So the 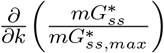 in Eq. (S12) is larger than 0, finishing our proof about why decreasing *k* results in the decrease of the 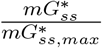.

As for the DoRA of tGTPase, the 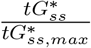 is also positively correlated with the mGTPase activation level if only kinetic parameter is varied. As we can see from the above paragraph, the kinetic parameters contributing to the mGTPase activation level appear in *F*_1_ and *a*_*H*_, and thus affect 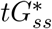 only though *F*_2_. First we studied how increasing *a*_*H*_, i.e., decreasing the mGTPase activation level, influences the 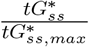. Similarly to the case of mGTPase, the sign of 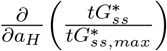 is determined by 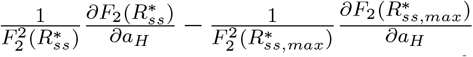 and 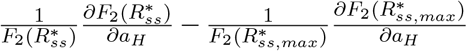. Combined with the definition of 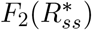, the 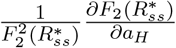 and 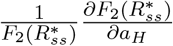 can be rewritten as:

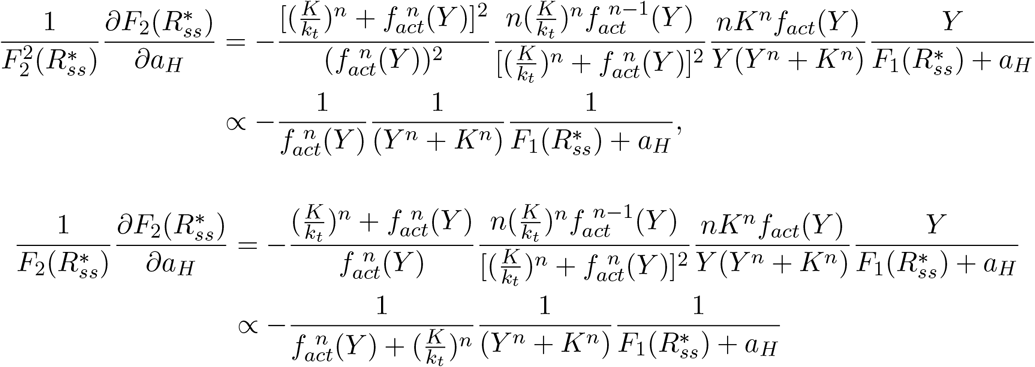

where 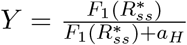 and 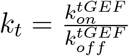. So 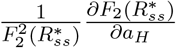 and 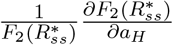 are both increasing function of 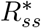, leading the negative sign of the derivative of 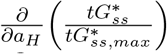 with respect to the *a*_*H*_. This indicates that the large *a*_*H*_ not only decreases the mGTPase activation level but also lowers the 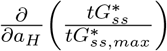. For the case where only *k* is varied, decreasing *k* causes both the low mGTPase activation level and the low 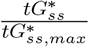, because 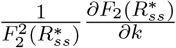 and 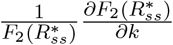 are both decreasing functions of 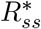 as shown below:

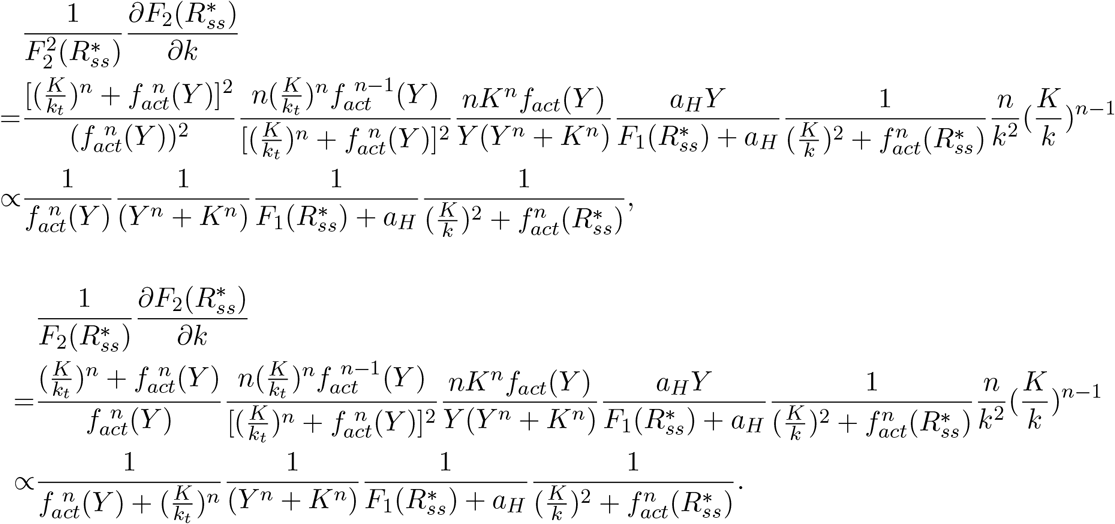

The above analyses showed that tuning one kinetic parameter to achieve low mGTPase activation level leads to the low 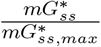 and 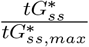. So, the 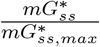 or 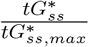 crosses the receptor curve from the left to the right. This observation suggests the DoRA metric experiences a decreasing and increasing trend (Figure 2C). It should be noted that we didn’t consider the effects of changing kinetic parameters *B* or *EC*_50_ in the function *f*_*act*_.

## 3 Analysis for the feedback’s effect on the DoRA

### 3.1 The mass action model

We first analyzed how the DoRA is affected by the negative feedback in the mass action model. The steady state of the mGAP concentration is given by

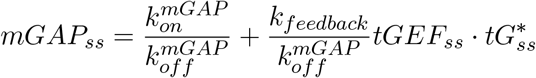

for the AND logic gate, and

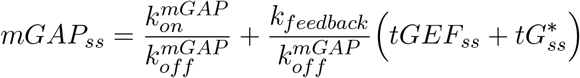

for the OR logic gate. Therefore, the steady state of the *mG*^*\**^ can be written as:

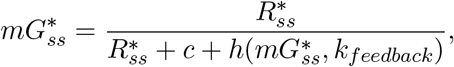

Where

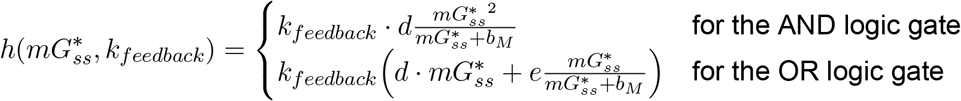

Here, *c, d, e, b*_*M*_ are constants and defined as follows: 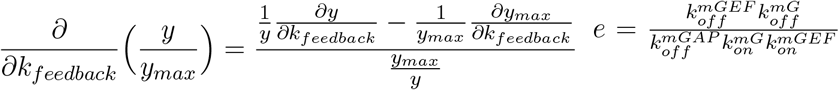, and 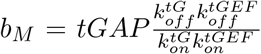. Note that the 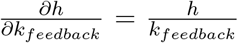 holds for any logic gate. For simplicity, we used *x* and *y* to denote the 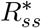 and 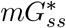, respectively. Then relation between 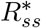 and 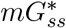 can be rewritten as:

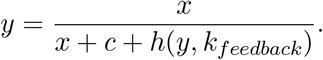

In order to know the normalized fractional activation change when increasing feedback strength, we used *y*_*max*_ representing the maximal value of *y* when *x* is 1, and derived the expression of 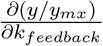 as follows:

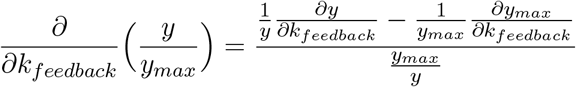

Since the denominator is positive, the sign of the 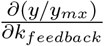 is only determined by the monotonicity of the function 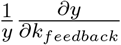. Then we rewrote this function as follows:

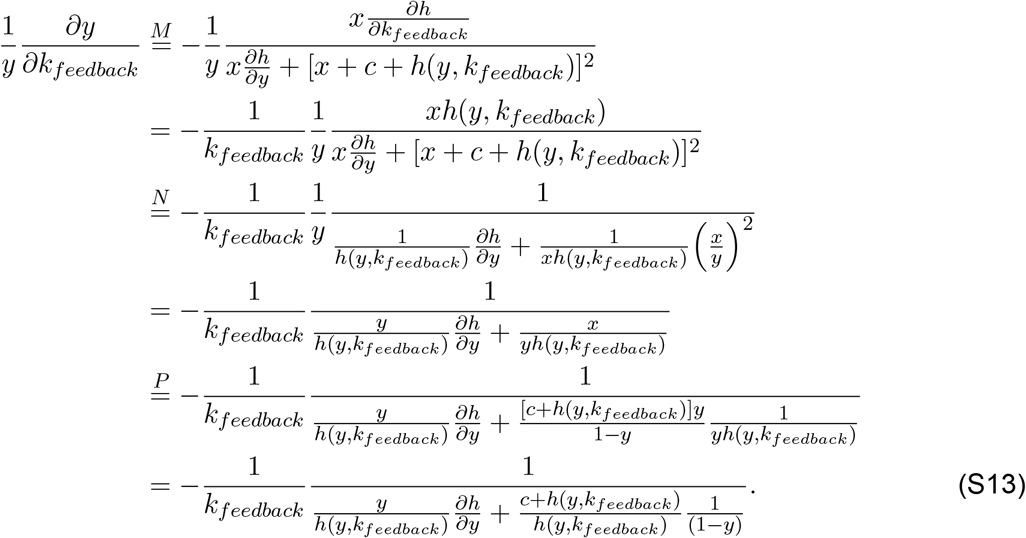

Here, the equation *M* is obtained because 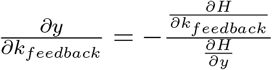 where 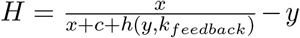. The equation *N* is based on the relation between *x* and *y*, i.e., 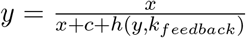. This relation can also be rewritten as 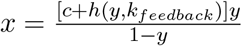, which leads to the equation *P*.

From the above analysis, we showed that the sign of the 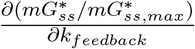 is determined by the difference of values of the function 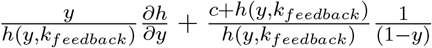 at 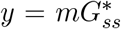 and 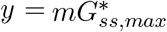. If this function at 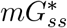 is smaller than that at 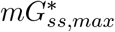, the sign of the 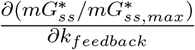 is negative; however, the larger value at 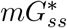 than that at 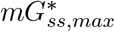 indicates the positive sign. Then we focused on the monotonicity of this function with respect to *y*. For simplicity, we used *h*(*y*) to represent *h*(*y, k*_*feedback*_). The first term 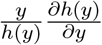, i.e., 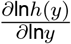, reflects the kinetic order: if *h*(*y*) is of second kinetic order in *y*, i.e., *h*(*y*) = *y*^2^, then 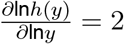. In fact, in our model, 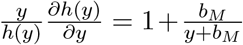 for the AND logic gate, so the first term is an increasing function all the time; but for the OR logic gate, 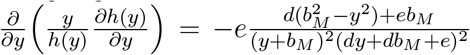, so the function 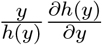 decreases and then increases if *y* is larger enough. But this function seems to be flat in most range of *y* (Figure S4-S7). The second term 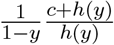 can be approximated by 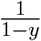 when *y* is large, because the basal production rate of mGAP 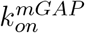 is small and thus the constant *c* can be neglected. Therefore, the second term is an increasing function when *y* is large. In fact, when *y* is small, the existence of the 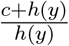 makes the whole function decrease with increased *y*. The rigorous derivation of the monotonicity of the second term is as follows: 1) the derivative of this function is 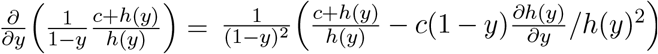; 2) the derivative of the function in the bracket is 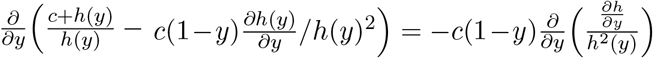; 3) 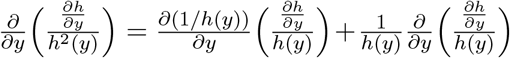 is smaller than 0, 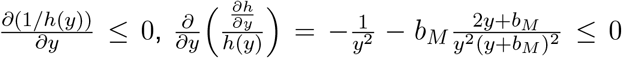 for the AND logic gate, and 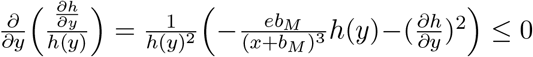 for the OR logic. 4) so the function 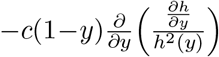 is always larger than 0, and thus the function 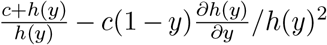 increases monotonically with respect to *y*; 5) combining facts that the function 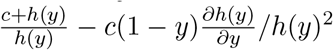 is smaller than 0 at *y* = 0 and larger than 0 when *y* = 1, this function is negative and then becomes positive with increased *y*; 6) as the sign of this function determines the trend of the function 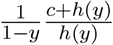, it can be concluded that the function 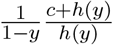 will decrease and then increase when *y* increases. Recall that the first term in 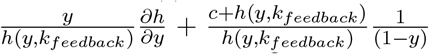 is almost flat, the whole function trend is mainly determined by the second term. So, if 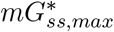 is large and located in the increasing branch, the function value at 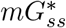 has a large probability to be smaller than that at 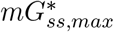, leading to the negative sign of 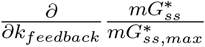. However, if 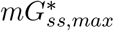 is very small which is located in the decreasing branch, the value of the function at 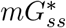 is larger than that at 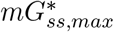, resulting the positive sign of the 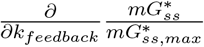.

In all, the high (or low) level fo mGTPase activation may lead to the descent (or ascent) of the 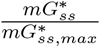 when increasing feedback strength. Based on the fact that 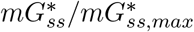 curve is always higher than the receptor curve (due to 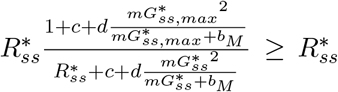), the descent and ascent of the 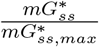 correspond to the improvement and impairment of the DoRA, respectively. Therefore, when mGTPase activation level is high, the feedback enhances the DoRA of mGTPase; such DoRA is impaired by the feedback if mGTPase activation level is low. We also compared the derivatives of the distance with respect to the feedback strength from numerical simulations and analytical analysis; they have a godd fit (Figure S4-S7). Up to now, we finished the validation of the effect of feedback on the DoRA of mGTPases.

Next we explored the role of the feedback on the DoRA of tGTPase. The steady state of *tG*^*\**^ is given by:

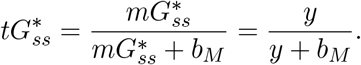

where the constant *b*_*M*_ has been defined in the previous section. The *y* is still used to denote 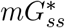. Similarly, we still focused on the 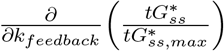, that is, 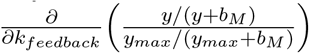. This function can be rewritten as:

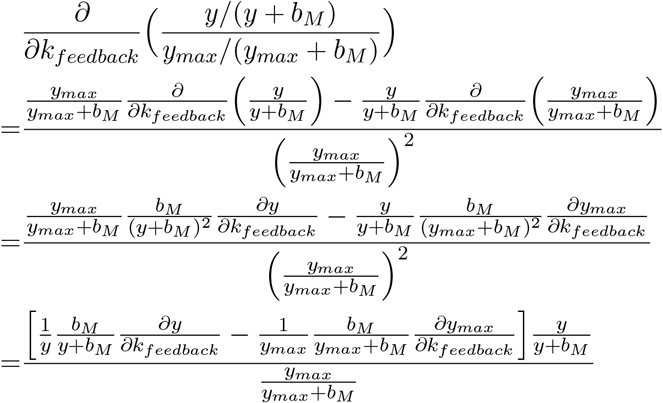

So the sign of the 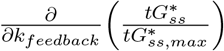 is determined by the difference of the values of the function 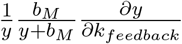 at 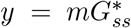 and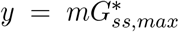. Based on our previous analysis about the 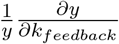, we 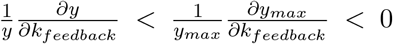 for the high mGTPase activation level and 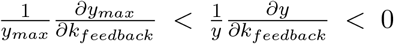 for the low mGTPase activation level. Therefore, for the high mGTPase activation level, the 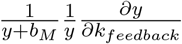 is smaller than 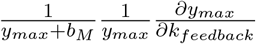 due to 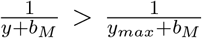; for the low mGTPase activation level, 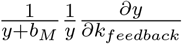 might maintain a larger value than 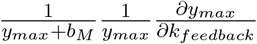. Besides, because 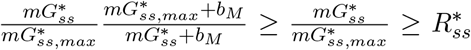, the 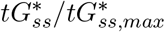 curve is always higher than the receptor curve. Taken together, the DoRA of tGTPase is like that of mGTPase, i.e., the high (or low) mGTPase activation level indicates the feedback’s positive (or negative) role on the DoRA.

### 3.2 The Hill-function model

In this section, we focused on the feedback’s role in the DoRA for the Hill-function model. The steady state of the mGAP concentration is given by

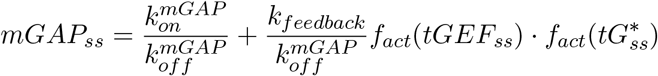

for the AND logic gate, and

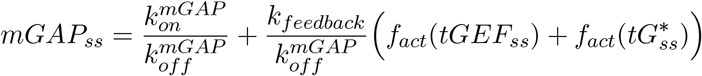

for the OR logic gate. By substituting the *mGAP*_*ss*_ with the above equations, we got the 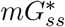 as follows:

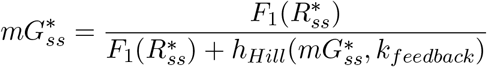

where 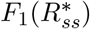 has been defined in the previous section. The 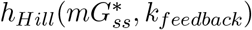 is defined as:

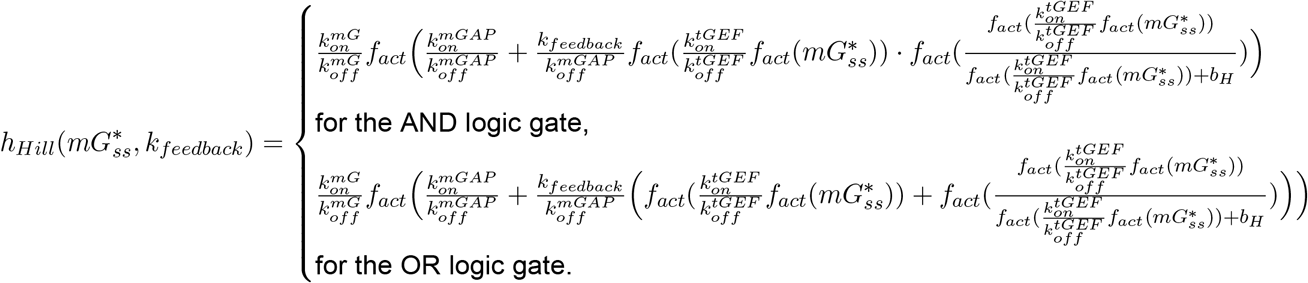

For simplicity, we still used *x* and *y* to denote the 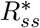 and 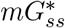, respectively. Then relation between 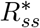 and 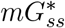 can be rewritten as:

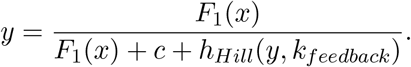

As we have shown in the previous section, the sign of the 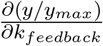 is determined 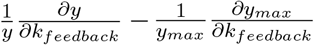. The expression of 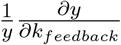 is shown as follows:

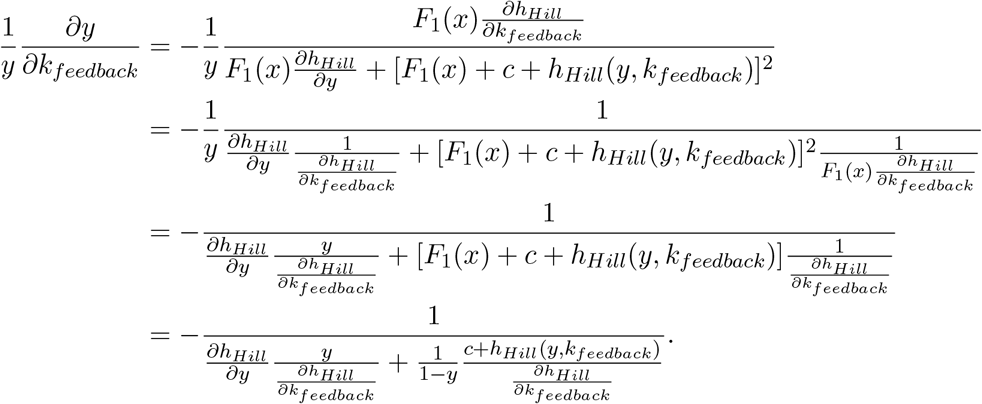

To further simplify this equation, we rewrote the *h*_*Hill*_ *(y)* as *G*(*z*), where 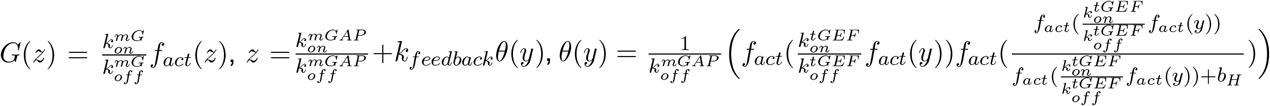 for the AND logic gate and 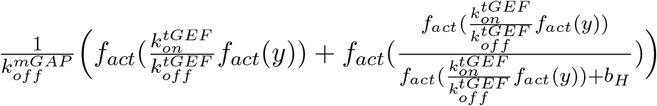 for the OR logic gate. Therefore, the derivative of the second term in the denominator with respect to the *y* is given by:

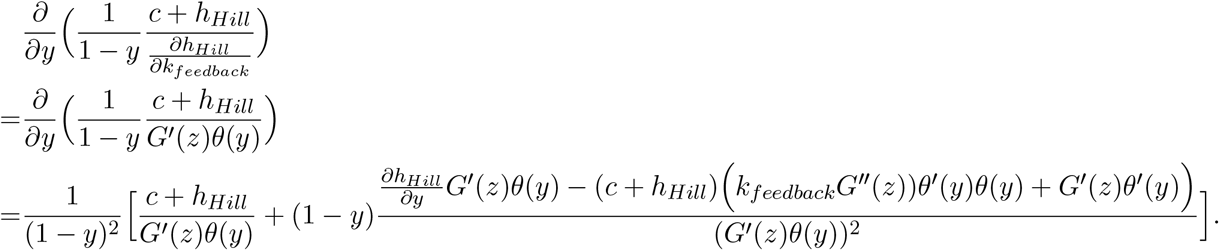

When *y* goes to 1, this equation is larger than 0. When *y* is 0, *θ*(0) = 0, and 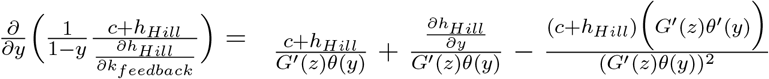; due to 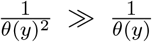 when *y* goes to 0, the third term dominates and thus 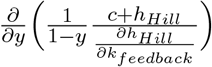 is smaller than 0. To now, we have proved that the second term in the denominator of the function 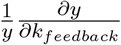 is decreasing for the small *y* and increasing for enough large *y*. If we neglected the effect of the first term, we can conclude that 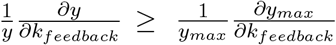 for the small *y*_*max*_ and 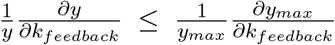 for the large *y*_*max*_. Since the large *y*_*max*_ usually corresponds to the case that the normalized mGTPase curve is higher than the receptor curve, the negative sign of the 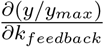 indicates the closer distance between two curves with increased feedback. Similarly, the small *y*_*max*_ also suggests the closer distance between two curves with increased feedback, because at that time the normalized mGTPase curve is usually lower than the receptor curve. To now, we have finished the proof of Figure 3C. The DoRA of tGTPase follows the same rule, and it can be proven using the same method shown in the previous section.

## 4. Analysis for different logic gate

### 4.1 The mass action model

In the mass action model, the OR logic gates show a little better DoRA behavior than AND logic gate. We used 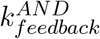 and 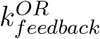 to represent the negative feedback strength for the system with the AND logic gate and the system with the OR logic gate, respectively. The requirement about the same *mG*^*\**^ activation level (i.e., 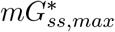) indicates the same mGAP level when the stimulus *S* → ∞, because the mGEF is not affected by logic gates. Since the mGAP level is the same, the AND and OR logic gates show the same regulatory strength to the mGAP, leading to the following equation:

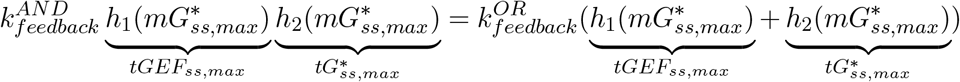

where 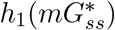 and 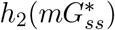 denote *tGEF*_*ss*_ and 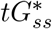, respectively. All variables are steady-state values when the stimulus *S* → ∞. Because the function 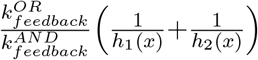 is a decreasing function of *x*, we had

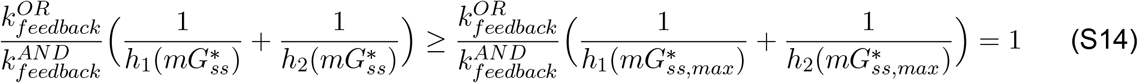

The left-hand side can be rewritten as 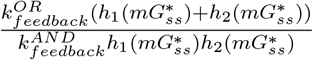. So, 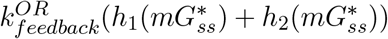 is larger than 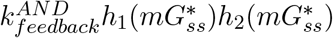; this indicates that the system with the OR logic gate has stronger feedback strength and thus lower *mG*^*\**^ level when compared to the system with the AND logic gate. Combined with the fact that 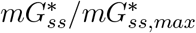 curve is always higher than the receptor curve in the mass action model, the distance between these two curves in the system with the OR logic gate is smaller than that with AND logic gate (Figure 4B). This implies the better DoRA performance of the OR logic in the mass action model. However, this advantage is small (See main texts for more explanations).

---

**Table S1:** Parameters used in numerical simulations

|  |  |
| --- | --- |
| Stimulus $S$ | $10^x$, where $x$ is uniformly distributed in $[-5, 3]$ with the increment 0.2 |
| $k_{on}^i, i = mGEF, tGEF$ | 1 |
| $k_{off}^i, i = R^*, mGEF, mG^*, tGEF, tG^*, mGAP$ | 1 |
| $tGAP$ | 0.5 |
| $K$ | 0.5 |
| n (unless otherwise specified) | 1.5 |
| **Parameters used in Figure 2** |  |
| $k_{feedback}$ | 0 |
| $k_{on}^{mGAP}$ | [0.01:0.02:0.1, 0.15:0.05:0.4, 0.5:0.1:1] |
| $(k_{on}^R, k_{on}^{mG}, k_{on}^{tG})$ in the upper panel in B | $(0.5, 10^0, 10^0)$ |
| $k_{on}^R, k_{on}^{mG}, k_{on}^{tG}$ in the middle and lower panels in B | $k_{on}^R = 0.5$.<br>$k_{on}^{mG}$ and $k_{on}^{tG}$ are $10^x$, where $x$ is uniformly distributed in $[-2, 2]$ with the increment 0.2 in the lower panel or 0.1 in the middle panel |
| $(k_{on}^R, k_{on}^{mG}, k_{on}^{tG})$ (orange) in the upper panel in D | $(10^{0.2}, 10^{0.2}, 1)$ |
| $(k_{on}^R, k_{on}^{mG}, k_{on}^{tG})$ (magenta) in the upper panel in D | $(10^{-0.8}, 10^0, 1)$ |
| $(k_{on}^R, k_{on}^{mG}, k_{on}^{tG})$ (purple) in the upper panel in D | $(10^{-1.2}, 10^{-1.2}, 1)$ |
| $k_{on}^R, k_{on}^{mG}, k_{on}^{tG}$ in the middle and lower panels in D | $k_{on}^R$ and $k_{on}^{mG}$ are $10^x$, where $x$ is uniformly distributed in $[-2, 2]$ with the increment 0.2.<br>$k_{on}^{tG} = 0.01, 1, \text{ or } 10$ |
| **Parameters used in Figure 3** |  |
| $k_{on}^{mGAP}$ | 0.5 |
| $k_{feedback}$ | 0 and $10^x$, $x$ is uniformly distributed in $[-1, 2.6]$ with the increment 0.2 |
| $(k_{on}^R, k_{on}^{mG}, k_{on}^{tG})$ (purple) in the upper two panels in B | $(0.5, 10^{-1.4}, 10^1)$ |
| $(k_{on}^R, k_{on}^{mG}, k_{on}^{tG})$ (orange) in the upper left panel in B | $(0.5, 10^{1.4}, 10^2)$ |
| $(k_{on}^R, k_{on}^{mG}, k_{on}^{tG})$ (magenta) in the upper left panel in B | $(0.5, 10^{0.4}, 10^1)$ |
| $(k_{on}^R, k_{on}^{mG}, k_{on}^{tG})$ (black) in the upper right panel in B | $(0.5, 10^{-0.6}, 10^{0.6})$ |
| $(k_{on}^R, k_{on}^{mG}, k_{on}^{tG})$ (orange) in the upper right panel in B | $(0.5, 10^1, 10^1)$ |
| $k_{on}^R, k_{on}^{mG}, k_{on}^{tG}$ in the middle and lower panels in B | Same as those in Figure 2B |
| $(k_{on}^R, k_{on}^{mG}, k_{on}^{tG})$ (orange in right upper corner) in upper panels in D | $(10^{0.4}, 10^{0.4}, 1)$ |
| $(k_{on}^R, k_{on}^{mG}, k_{on}^{tG})$ (magenta) in upper panels in D | $(10^{-1}, 10^1, 1)$ |
| $(k_{on}^R, k_{on}^{mG}, k_{on}^{tG})$ (orange in left bottom corner) in upper panels in D | $(10^{-0.6}, 10^{-0.6}, 1)$ |
| $(k_{on}^R, k_{on}^{mG}, k_{on}^{tG})$ (black) in upper panels in D | $(10^1, 10^{-0.6}, 1)$ |
| $k_{on}^R, k_{on}^{mG}, k_{on}^{tG}$ in the middle and lower panels in D | Same as those in Figure 2D |
| **Parameters used in Figure S4 and Figure S6** |  |
| $k_{on}^{mGAP}$ | 0.5 |
| $(k_{on}^R, k_{on}^{mG}, k_{on}^{tG})$ | (1, 10, 10) |
| $k_{feedback}$ | 1, 5, or 10 |
| **Parameters used in Figure S5 and Figure S7** |  |
| $k_{on}^{mGAP}$ | 0.5 |
| $(k_{on}^R, k_{on}^{mG}, k_{on}^{tG})$ | $(0.1, 10^{-0.5}, 10^{-0.5})$ |
| $k_{feedback}$ | 1, 5, or 10 |

**Figure S1:**
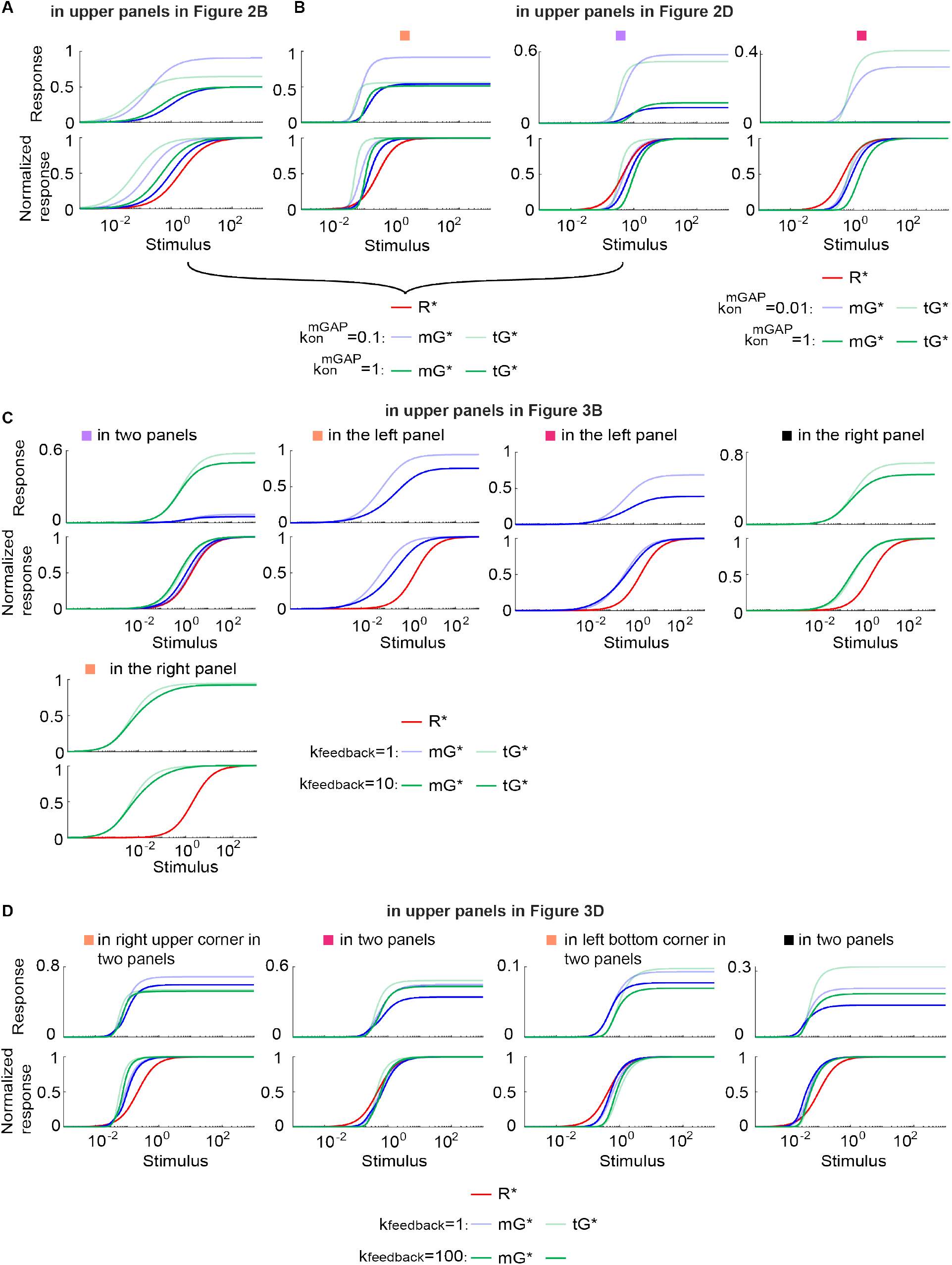
The dose-response curves for the parameter sets in Table S1.

**Figure S2:**
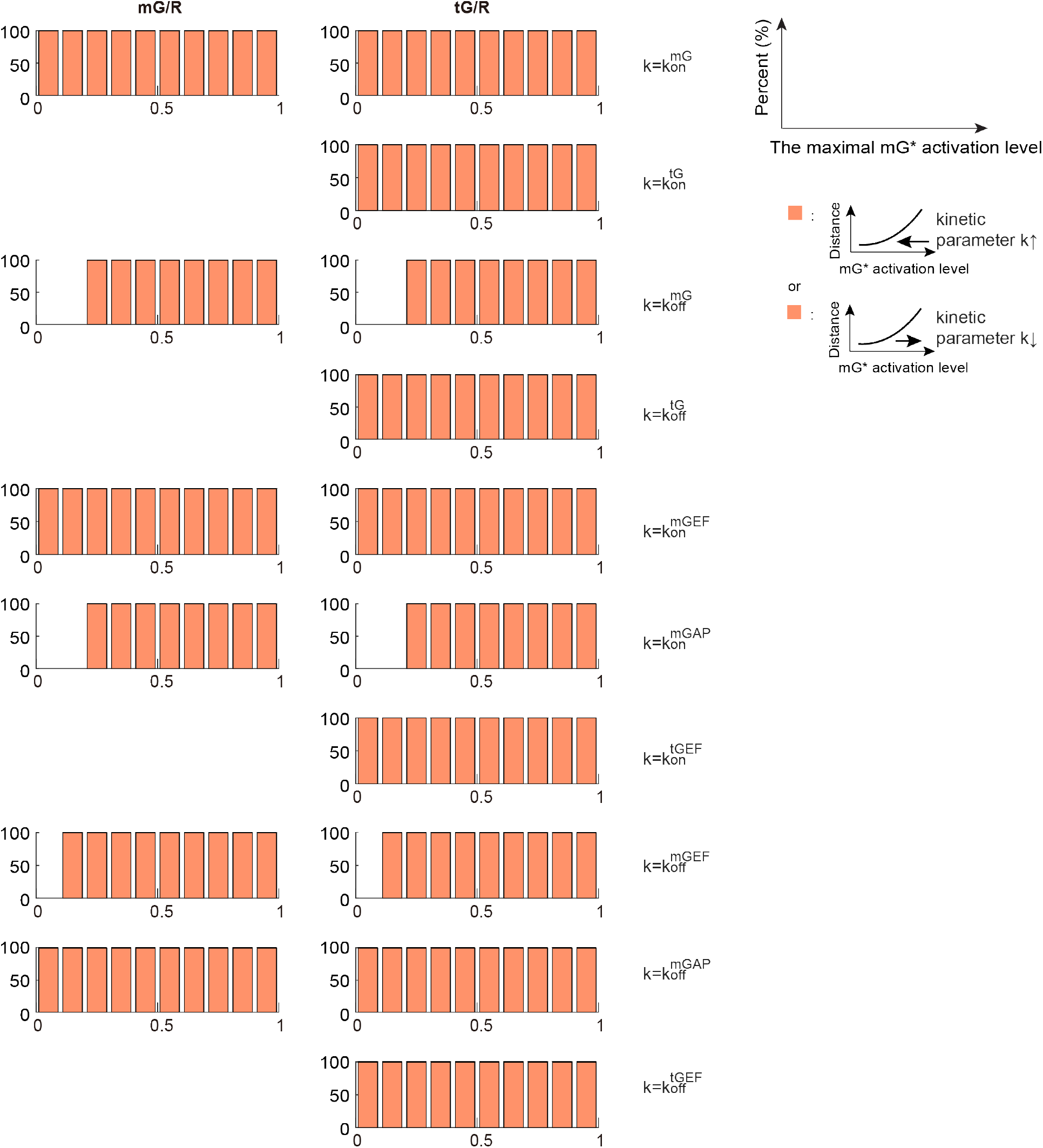
The distributions of trends for every kinetic parameter in coupled switches without feedback when using the mass action model. Same plots as in the middle panel Figure 2B except that the varied kinetic parameter is different. The varied kinetic parameter is shown on the right. Some of the left panels are missing, because this kinetic parameter cannot affect the DoRA metric for mGTPase. One thousand parameter sets are randomly assigned in a logarithmic scale in the whole parameter space using Latin hypercubic sampling, and each parameter is in the range [10^*−*2^, 10^1^]. Then the varied parameter is uniformly changed from 10^*−*2^ to 10^1^ in a logarithmic scale.

**Figure S3:**
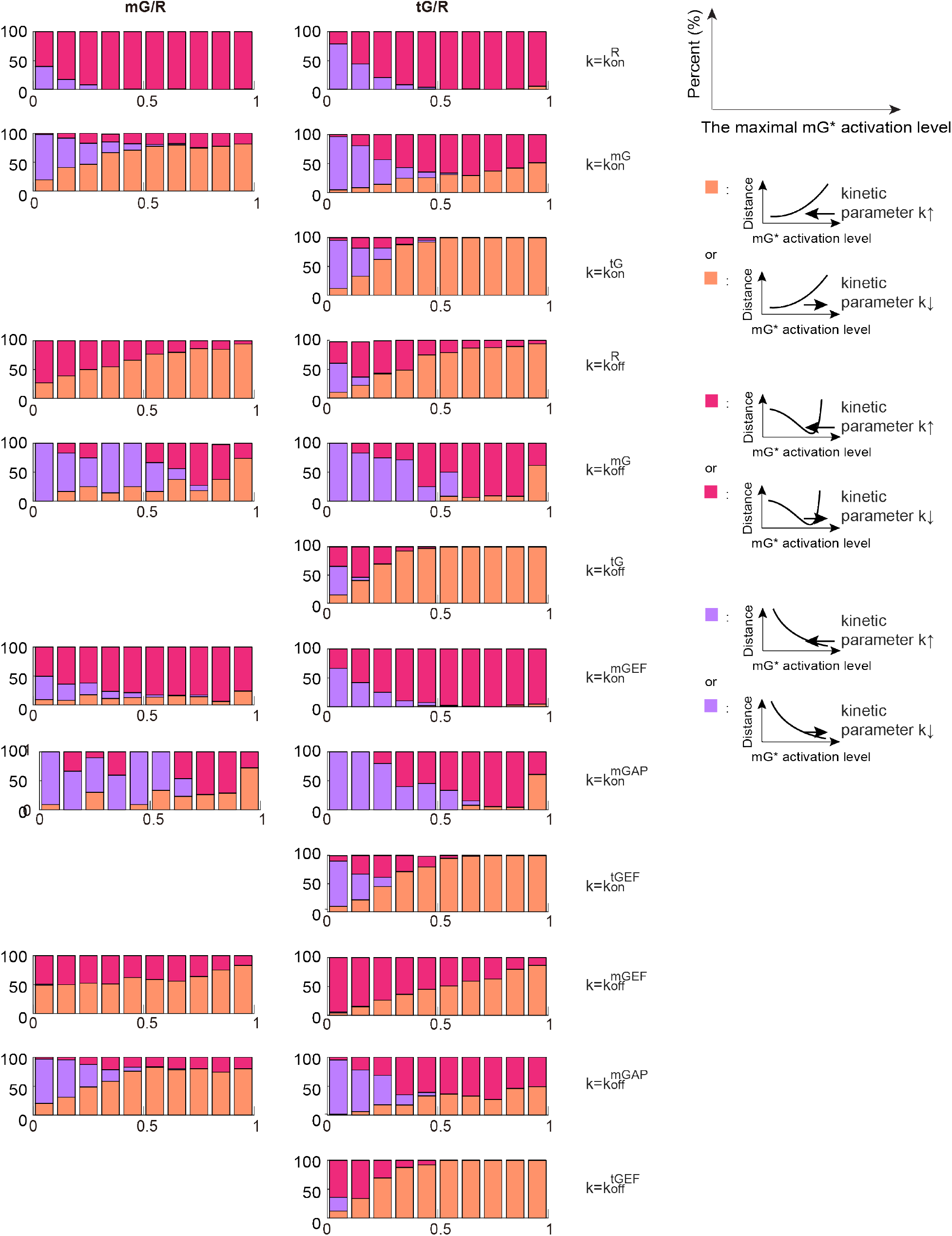
Same plot as Figure S2 except that the Hill-function kinetics is adopted.

**Figure S4:**
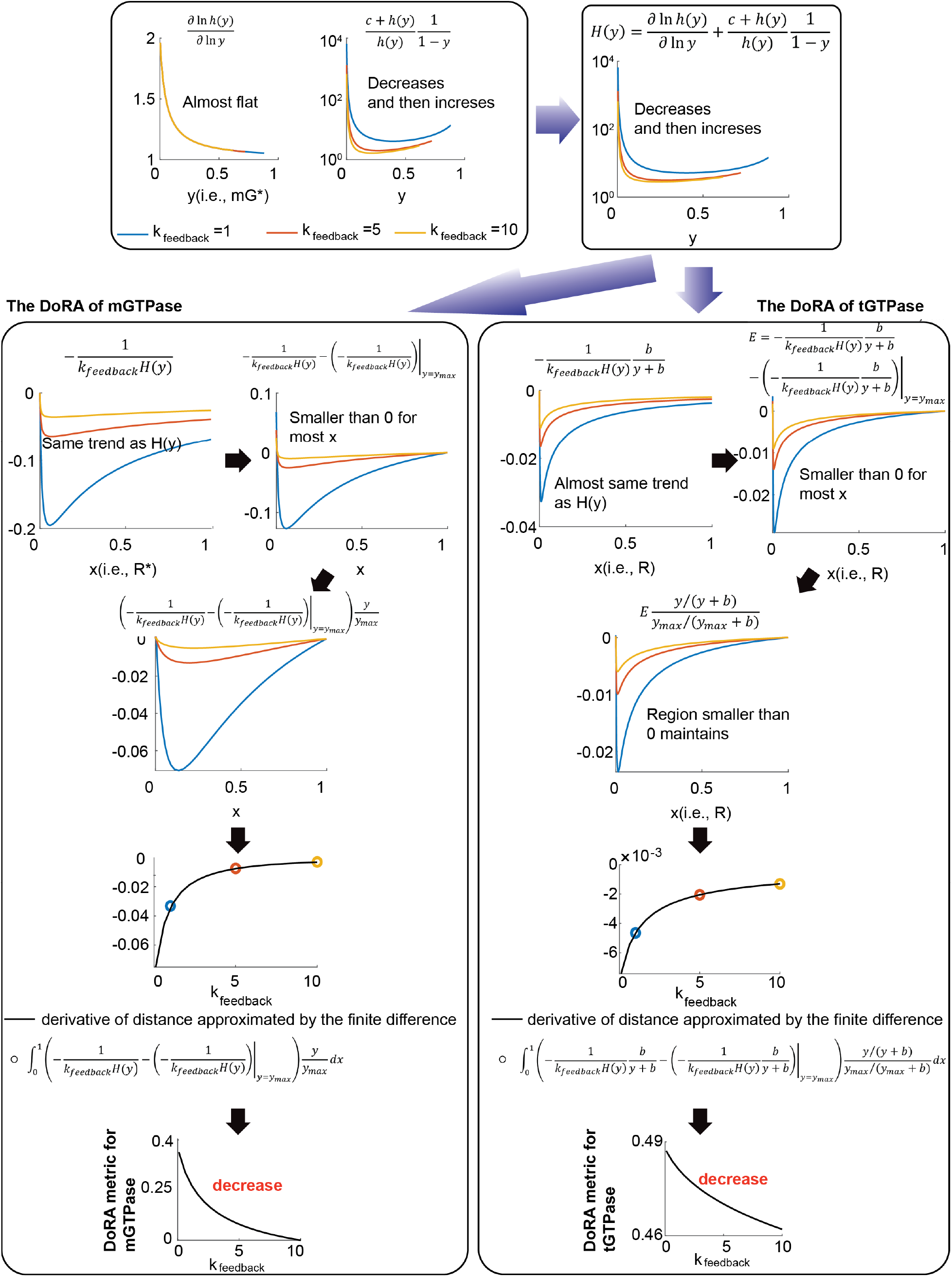
DoRA of the m- and tGTPase is improved with increased feedback strength in the mass action model. Here AND gate is applied. See Table S1 for parameters.

**Figure S5:**
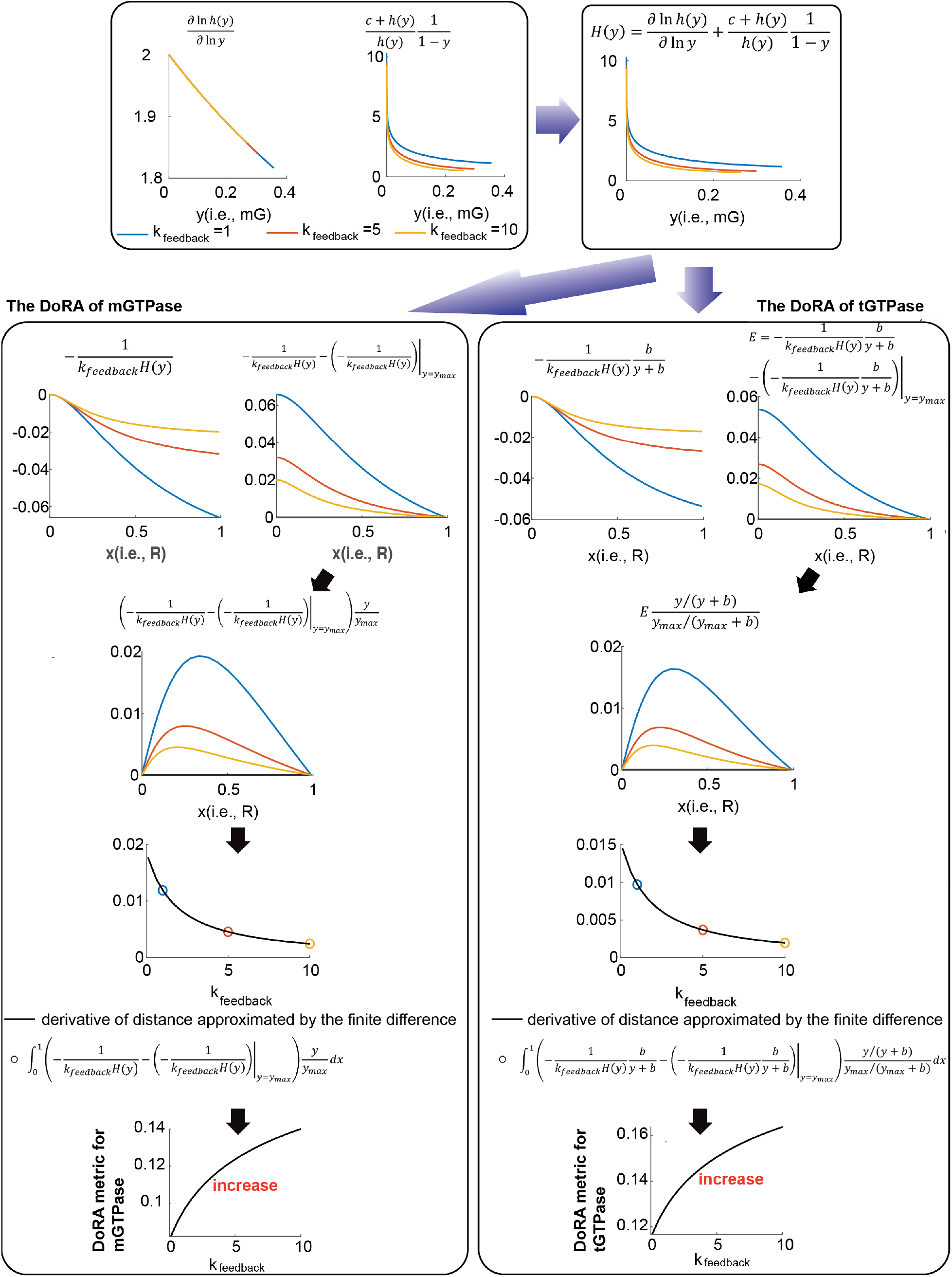
For some kinetic parameter sets, DoRA is impaired by increased feedback strength in the mass action model. Here AND gate is applied. See Table S1 for parameters.

**Figure S6:**
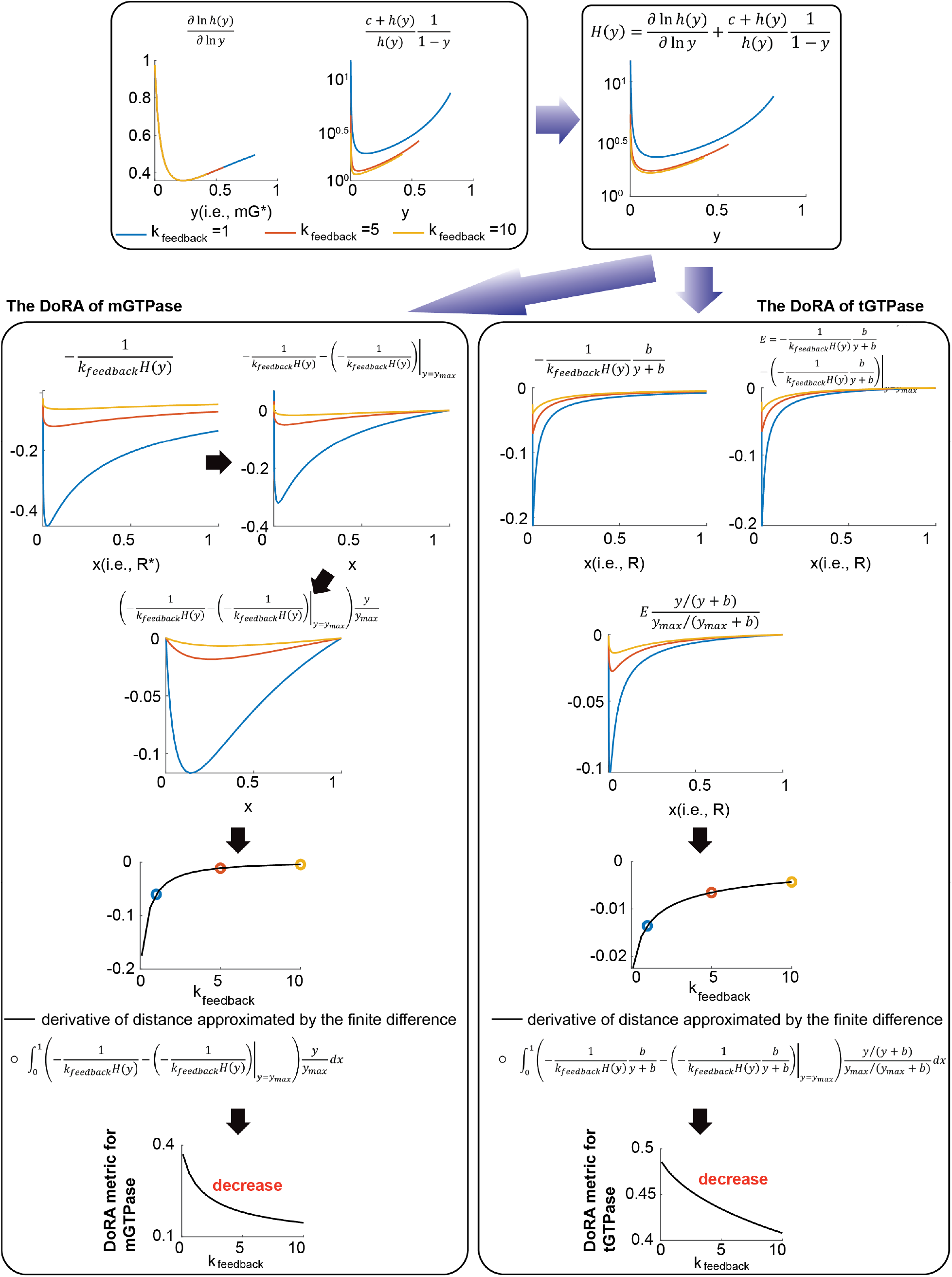
Same plot as Figure S4 but OR logic is used to model the negative feedback. See Table S1 for parameters.

**Figure S7:**
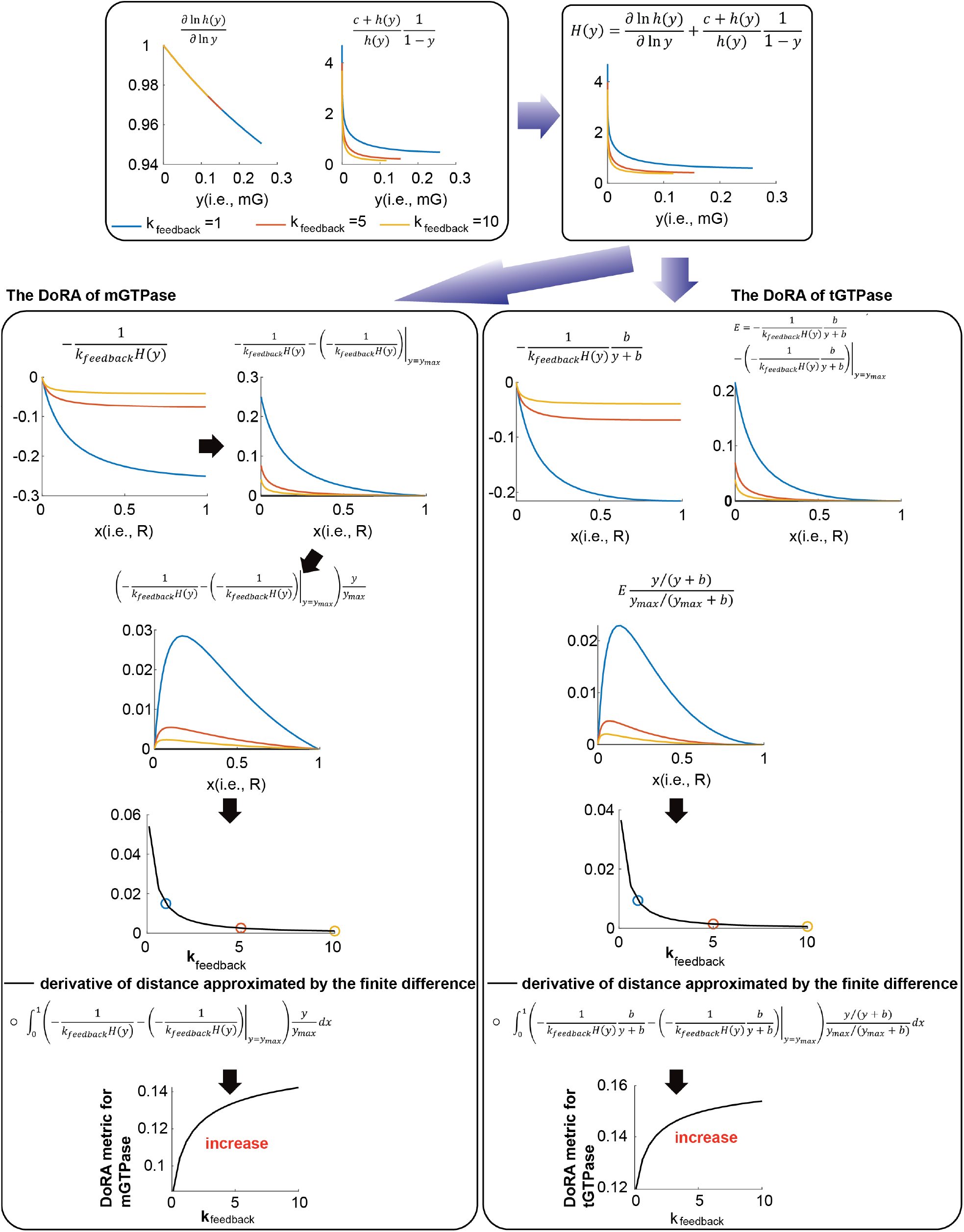
Same plot as Figure S5 but OR logic is used to model the negative feedback. See Table S1 for parameters.

**Figure S8:**
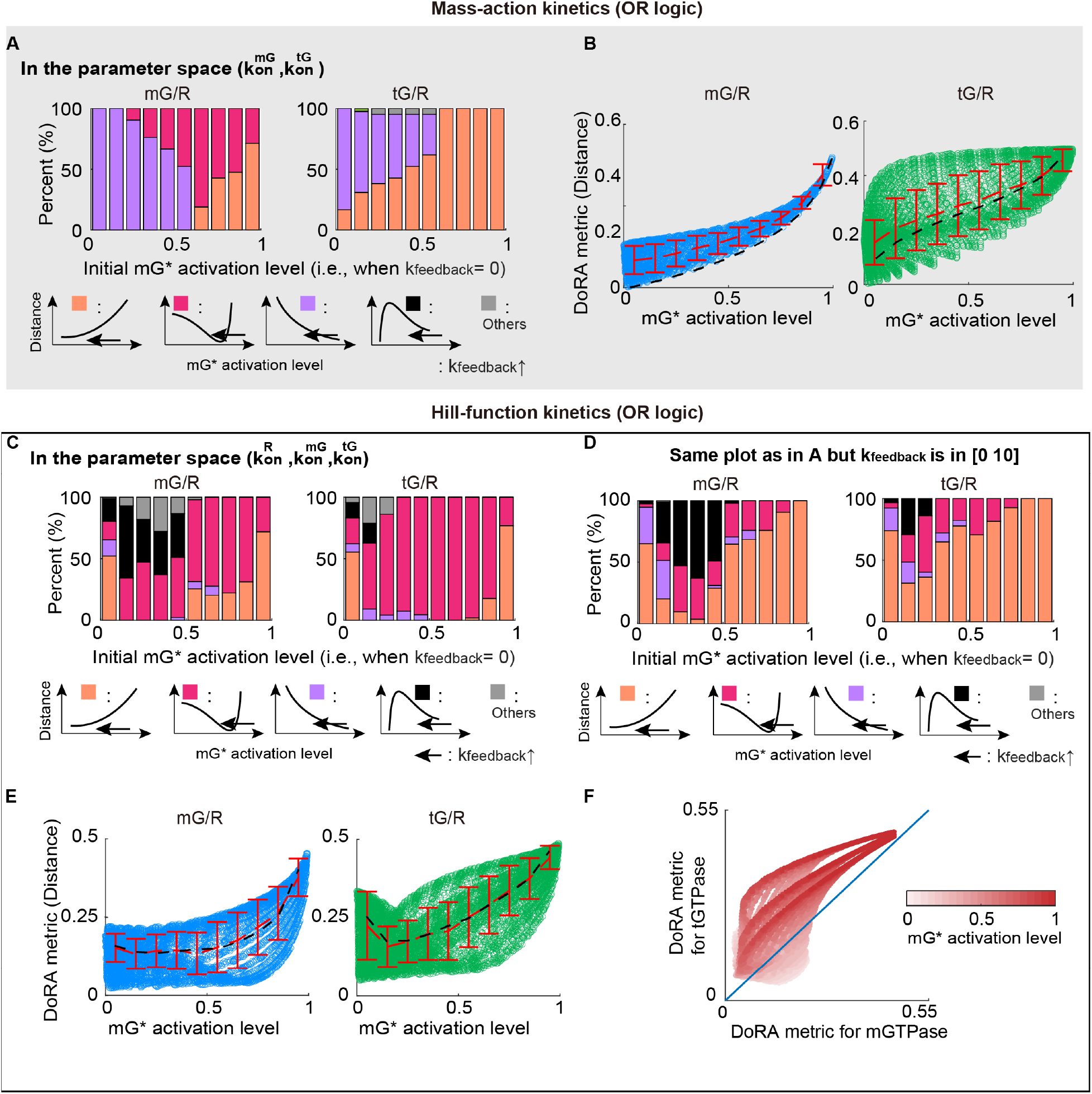
The simulation results when using OR logic gate. (A-B) Same plot as the middle and bottom panels in Figure 3B except that OR logic is used to model the negative feedback. (C-E) Same plot as the middle and bottom panels in Figure 3D except the logic gate. The panel C and D has different ranges of *k*_*feedback*_. (F) Same plot as the right panel in Figure 5D except the logic gate.

**Figure S9:**
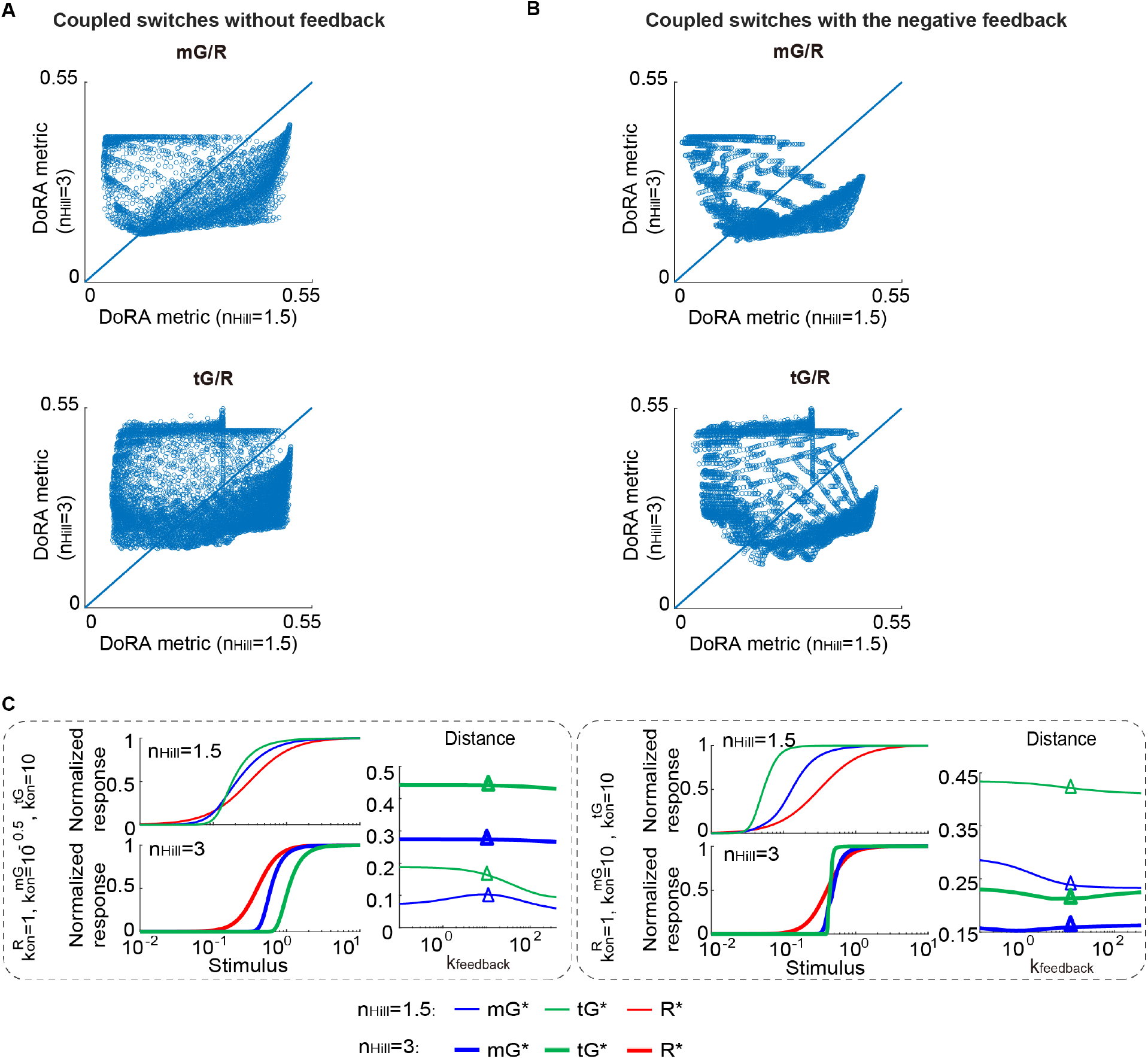
The effect of the Hill coefficient. (A-B) The scatter plots of the DoRA metric with *n*_*Hill*_ 1.5 versus that with *n*_*Hill*_ 3 in the parameter space 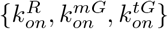 for the circuit without (A) or with feedback (B). The *n*_*Hill*_ is the Hill coefficient *n* mentioned in the first section. (C) Two examples showing the different effects of *n*_*Hill*_ on DoRA.

## References

1. H. G. Dohlman, J. Thorner, Regulation of G Protein–Initiated Signal Transduction in Yeast: Paradigms and Principles. Annual Review of Biochemistry 70, PMID: 11395421, 703–754, DOI 10.1146/annurev.biochem.70.1.703, eprint: https://doi.org/10.1146/annurev.biochem.70.1.703, (https://doi.org/10.1146/annurev.biochem.70.1.703) (2001).

2. R. Sando, T. C. Südhof, Latrophilin GPCR signaling mediates synapse formation. eLife 10, ed. by G. W. Davis, G. L. Westbrook, G. W. Davis, e65717, ISSN: 2050-084X, DOI 10.7554/eLife.65717, (https://doi.org/10.7554/eLife.65717) (Mar. 2021).

3. K. K. Midde et al., Multimodular biosensors reveal a novel platform for activation of G proteins by growth factor receptors. Proceedings of the National Academy of Sciences 112, E937–E946, DOI 10.1073/pnas.1420140112, eprint: https://www.pnas.org/doi/pdf/10.1073/pnas.1420140112, (https://www.pnas.org/doi/abs/10.1073/pnas.1420140112) (2015).

4. R. C. Yu et al., Negative feedback that improves information transmission in yeast signalling. Nature 456, 755–761, ISSN: 1476-4687, DOI 10.1038/nature07513, (https://doi.org/10.1038/nature07513) (Dec. 2008).

5. X. Chen, M. D. Resh, Cholesterol Depletion from the Plasma Membrane Triggers Ligand independent Activation of the Epidermal Growth Factor Receptor *. Journal of Biological Chemistry 277, 49631–49637, ISSN: 0021-9258, DOI 10.1074/jbc.M208327200, (https://doi.org/10.1074/jbc.M208327200) (Dec. 2002).

6. G. Shinar, R. Milo, M. R. Martínez, U. Alon, Input–output robustness in simple bacterial signaling systems. Proceedings of the National Academy of Sciences 104, 19931–19935, ISSN: 0027-8424, DOI 10.1073/pnas.0706792104, eprint: https://www.pnas.org/content/104/50/19931.full.pdf, (https://www.pnas.org/content/104/50/19931) (2007).

7. K. R. Ghusinga, R. D. Jones, A. M. Jones, T. C. Elston, Molecular switch architecture determines response properties of signaling pathways. Proceedings of the National Academy of Sciences 118, ISSN: 0027-8424, DOI 10.1073/pnas.2013401118, eprint: https://www.pnas.org/content/118/11/e2013401118.full.pdf, (https://www.pnas.org/content/118/11/e2013401118) (2021).

8. L. Yan, Q. Ouyang, H. Wang, DoseResponse Aligned Circuits in Signaling Systems. PLOS ONE 7, 1–10, DOI 10.1371/journal.pone.0034727, (https://doi.org/10.1371/journal.pone.0034727) (Apr. 2012).

9. S. S. Andrews, W. J. Peria, R. C. Yu, A. ColmanLerner, R. Brent, PushPull and Feedback Mechanisms Can Align Signaling System Outputs with Inputs. Cell Systems 3, 444–455.e2, ISSN: 24054712, DOI https://doi.org/10.1016/j.cels.2016.10.002, (https://www.sciencedirect.com/science/article/pii/S2405471216303210) (2016).

10. M. Adler, A. Mayo, U. Alon, Logarithmic and Power Law InputOutput Relations in Sensory Systems with FoldChange Detection. PLOS Computational Biology 10, 1–14, DOI 10.1371/journal.pcbi.1003781, (https://doi.org/10.1371/journal.pcbi.1003781) (Aug. 2014).

11. J. Paulsson, Summing up the noise in gene networks. Nature 427, 415–418, ISSN: 1476-4687, DOI 10.1038/nature02257, (https://doi.org/10.1038/nature02257) (Jan. 2004).

12. T. Shibata, K. Fujimoto, Noisy signal amplification in ultrasensitive signal transduction. Proceedings of the National 102, 331–336, ISSN: 0027-8424, DOI 10.1073/pnas.0403350102, eprint: https://www.pnas.org/content/102/2/331.full.pdf, (https://www.pnas.org/content/102/2/331) (2005).

13. L. Wang, J. Xin, Q. Nie, A Critical Quantity for Noise Attenuation in Feedback Systems. PLOS Computational Biology 6, 1–17, DOI 10.1371/journal.pcbi.1000764, (https://doi.org/10.1371/journal.pcbi.1000764) (Apr. 2010).

14. L. PotvinTrottier, N. D. Lord, G. Vinnicombe, J. Paulsson, Synchronous longterm oscilla tions in a synthetic gene circuit. Nature 538, 514–517, ISSN: 1476-4687, DOI 10.1038/nature19841, (https://doi.org/10.1038/nature19841) (Oct. 2016).

15. G. Hennequin, Y. Ahmadian, D. B. Rubin, M. Lengyel, K. D. Miller, The Dynamical Regime of Sensory Cortex: Stable Dynamics around a Single StimulusTuned Attractor Account for Patterns of Noise Variability. Neuron 98, 846–860.e5, ISSN: 08966273, DOI https://doi.org/10.1016/j.neuron.2018.04.017, (https://www.sciencedirect.com/science/article/pii/S0896627318303258) (2018).

16. L. Qiao, W. Zhao, C. Tang, Q. Nie, L. Zhang, Network Topologies That Can Achieve Dual Function of Adaptation and Noise Attenuation. Cell Systems 9, 271–285.e7, ISSN: 2405 4712, DOI https://doi.org/10.1016/j.cels.2019.08.006, (https://www.sciencedirect.com/science/article/pii/S2405471219302753) (2019).

17. J. Selimkhanov et al., Accurate information transmission through dynamic biochemical signaling networks. Science 346, 1370–1373, DOI 10.1126/science.1254933, eprint: https://www.science.org/doi/pdf/10.1126/science.1254933, (https://www.science.org/doi/abs/10.1126/science.1254933) (2014).

18. R. Cheong, A. Rhee, C. J. Wang, I. Nemenman, A. Levchenko, Information Transduction Capacity of Noisy Biochemical Signaling Networks. Science 334, 354–358, DOI 10.1126/science.1204553, eprint: https://www.science.org/doi/pdf/10.1126/science.1204553, (https://www.science.org/doi/abs/10.1126/science.1204553) (2011).

19. S. Uda et al., Robustness and Compensation of Information Transmission of Signaling Pathways. Science 341, 558–561, DOI 10.1126/science.1234511, eprint: https://www.science.org/doi/pdf/10.1126/science.1234511, (https://www.science.org/doi/abs/10.1126/science.1234511) (2013).

20. S. Uda, S. Kuroda, Analysis of cellular signal transduction from an information theoretic approach. Seminars in Cell Developmental Biology 51, Information Theory in Systems Biology Xenopus as a model system for vertebrate development, 24–31, ISSN: 10849521, DOI https://doi.org/10.1016/j.semcdb.2015.12.011, (https://www.sciencedirect.com/science/article/pii/S1084952115300240) (2016).

21. P. Mehta, S. Goyal, T. Long, B. L. Bassler, N. S. Wingreen, Information processing and signal integration in bacterial quorum sensing. Molecular Systems Biology 5, 325, DOI https://doi.org/10.1038/msb.2009.79, eprint:https://www.embopress.org/doi/pdf/10.1038/msb.2009.79, (https://www.embopress.org/doi/abs/10.1038/msb.2009.79) (2009).

22. P. Cuatrecasas, InsulinReceptor Interactions in Adipose Tissue Cells: Direct Measurement and Properties. Proceedings of the National Academy of Sciences 68, 1264–1268, ISSN: 0027-8424, DOI 10.1073/pnas.68.6.1264, eprint: https://www.pnas.org/content/68/6/1264.full.pdf, (https://www.pnas.org/content/68/6/1264) (1971).

23. S. M. Amir, T. F. Carraway Jr., L. D. Kohn, R. J. Winand, The Binding of Thyrotropin to Isolated Bovine Thyroid Plasma Membranes. Journal of Biological Chemistry 248, 4092–4100, ISSN: 0021-9258, DOI 10.1016/S0021-9258(19)43843-X, (https://doi.org/10.1016/S0021-9258(19)43843-X) (June 1973).

24. S. Lin, T. Goodfriend, Angiotensin receptors. American Journal of PhysiologyLegacy Content 218, PMID: 4314569, 1319–1328, DOI 10.1152/ajplegacy.1970.218.5.1319, eprint: https://doi.org/10.1152/ajplegacy.1970.218.5.1319, (https://doi.org/10.1152/ajplegacy.1970.218.5.1319) (1970).

25. T. Nagashima et al., Quantitative Transcriptional Control of ErbB Receptor Signaling Under goes Graded to Biphasic Response for Cell Differentiation *<sup></sup>. Journal of Biological Chemistry 282, 4045–4056, ISSN: 0021-9258, DOI 10.1074/jbc.M608653200, (https://doi.org/10.1074/jbc.M608653200) (Feb. 2007).

26. H. Nunns, L. Goentoro, Signaling pathways as linear transmitters. eLife 7, ed. by W. Shou, A. Regev, W. Shou, S. S. Andrews, e33617, ISSN: 2050-084X, DOI 10.7554/eLife.33617, (https://doi.org/10.7554/eLife.33617) (Sept. 2018).

27. A. Bush et al., Yeast GPCR signaling reflects the fraction of occupied receptors, not the number. Molecular Systems Biology 12, 898, DOI https://doi.org/10.15252/msb.20166910, eprint:https://www.embopress.org/doi/pdf/10.15252/msb.20166910, (https://www.embopress.org/doi/abs/10.15252/msb.20166910) (2016).

28. U. Blank, G. Karlsson, S. Karlsson, Signaling pathways governing stemcell fate. Blood 111, 492–503, ISSN: 00064971, DOI https://doi.org/10.1182/blood-2007-07-075168, (https://www.sciencedirect.com/science/article/pii/S0006497120484629) (2008).

29. R. C. Sears, J. R. Nevins, Signaling Networks That Link Cell Proliferation and Cell Fate *. Journal of Biological Chemistry 277, 11617–11620, ISSN: 0021-9258, DOI 10.1074/jbc.R100063200, (https://doi.org/10.1074/jbc.R100063200) (Apr. 2002).

30. S. H. Patel, F. D. Camargo, D. Yimlamai, Hippo Signaling in the Liver Regulates Organ Size, Cell Fate, and Carcinogenesis. Gastroenterology 152, 533–545, ISSN: 00165085, DOI https://doi.org/10.1053/j.gastro.2016.10.047, (https://www.sciencedirect.com/science/article/pii/S0016508516355056) (2017).

31. K. Kessenbrock et al., A Role for Matrix Metalloproteinases in Regulating Mammary Stem Cell Function via the Wnt Signaling Pathway. Cell Stem Cell 13, 300–313, ISSN: 19345909, DOI https://doi.org/10.1016/j.stem.2013.06.005, (https://www.sciencedirect.com/science/article/pii/S1934590913002634) (2013).

32. M. S. Islam et al., Role of ActivinA and Myostatin and Their Signaling Pathway in Human Myometrial and Leiomyoma Cell Function. The Journal of Clinical Endocrinology Metabolism 99, E775–E785, ISSN: 0021-972X, DOI 10.1210/jc.2013-2623, eprint: https://academic.oup.com/jcem/article-pdf/99/5/E775/9050482/jcemE775.pdf, (https://doi.org/10.1210/jc.2013-2623) (May 2014).

33. I.C. Lo et al., Activation of Gαi at the Golgi by GIV/Girdin Imposes Finiteness in Arf1 Signaling. Developmental Cell 33, 189–203, ISSN: 15345807, DOI https://doi.org/10.1016/j.devcel.2015.02.009, (https://www.sciencedirect.com/science/article/pii/S1534580715001112) (2015).

34. O. Brandman, T. Meyer, Feedback Loops Shape Cellular Signals in Space and Time. Science 322, 390–395, DOI 10.1126/science.1160617, eprint: https://www.science.org/doi/pdf/10.1126/science.1160617, (https://www.science.org/doi/abs/10.1126/science.1160617) (2008).

35. M. Serrano, G. J. Hannon, D. Beach, A new regulatory motif in cellcycle control causing specific inhibition of cyclin D/CDK4. Nature 366, 704–707, ISSN: 1476-4687, DOI 10.1038/366704a0, (https://doi.org/10.1038/366704a0) (Dec. 1993).

36. I. Amit et al., A module of negative feedback regulators defines growth factor signaling. Nature Genetics 39, 503–512, ISSN: 1546-1718, DOI 10.1038/ng1987, (https://doi.org/10.1038/ng1987) (Apr. 2007).

37. S. Mangan, U. Alon, Structure and function of the feedforward loop network motif. Proceedings of the National 100, 11980–11985, DOI 10.1073/pnas.2133841100, eprint: https://www.pnas.org/doi/pdf/10.1073/pnas.2133841100, (https://www.pnas.org/doi/abs/10.1073/pnas.2133841100) (2003).

38. U. Alon, Network motifs: theory and experimental approaches. Nature Reviews Genetics 8, 450–461, ISSN: 1471-0064, DOI 10.1038/nrg2102, (https://doi.org/10.1038/nrg2102) (June 2007).

39. M. GarciaMarcos, P. Ghosh, M. G. Farquhar, GIV is a nonreceptor GEF for Gαi with a unique motif that regulates Akt signaling. Proceedings of the National Academy of Sciences 106, 3178–3183, ISSN: 0027-8424, DOI 10.1073/pnas.0900294106, eprint: https://www.pnas.org/content/106/9/3178.full.pdf, (https://www.pnas.org/content/106/9/3178) (2009).

40. N. A. Kalogriopoulos et al., Structural basis for GPCRindependent activation of heterotrimeric Gi proteins. Proceedings of the National Academy of Sciences 116, 16394–16403, ISSN: 0027-8424, DOI 10.1073/pnas.1906658116, eprint: https://www.pnas.org/content/116/33/16394.full.pdf, (https://www.pnas.org/content/116/33/16394) (2019).

41. N. Aznar, N. Kalogriopoulos, K. K. Midde, P. Ghosh, Heterotrimeric G protein signaling via GIV/Girdin: Breaking the rules of engagement, space, and time. BioEssays 38, 379–393, DOI https://doi.org/10.1002/bies.201500133, eprint:https://onlinelibrary.wiley.com/doi/pdf/10.1002/bies.201500133, (https://onlinelibrary.wiley.com/doi/abs/10.1002/bies.201500133) (2016).

42. P. Ghosh, M. GarciaMarcos, Do All Roads Lead to Rome in GProtein Activation? Trends in Biochemical Sciences 45, 182–184, ISSN: 09680004, DOI https://doi.org/10.1016/j.tibs.2019.10.010, (https://www.sciencedirect.com/science/article/pii/S0968000419302117) (2020).

43. L. M. Stolerman, P. Ghosh, P. Rangamani, Stability Analysis of a Signaling Circuit with Dual Species of GTPase Switches. Bulletin of Mathematical Biology 83, 34, ISSN: 1522-9602, DOI 10.1007/s11538-021-00864-w, (https://doi.org/10.1007/s11538-021-00864-w) (Feb. 2021).

44. L. Qiao et al., A Eukaryotic Circuit for SecreteandSense Autonomy. bioRxiv, DOI 10.1101/2021.03.18.436048, eprint: https://www.biorxiv.org/content/early/2021/10/03/2021.03.18.436048.full.pdf, (https://www.biorxiv.org/content/early/2021/10/03/2021.03.18.436048) (2021).

45. O. E. Sturm et al., The Mammalian MAPK/ERK Pathway Exhibits Properties of a Negative Feedback Amplifier. Science Signaling 3, ra90–ra90, DOI 10.1126/scisignal.2001212, eprint: https://www.science.org/doi/pdf/10.1126/scisignal.2001212, (https://www.science.org/doi/abs/10.1126/scisignal.2001212) (2010).

46. U. Bezeljak, H. Loya, B. Kaczmarek, T. E. Saunders, M. Loose, Stochastic activation and bistability in a Rab GTPase regulatory network. Proceedings of the National Academy of Sciences 117, 6540–6549, DOI 10.1073/pnas.1921027117, eprint: https://www.pnas.org/doi/pdf/10.1073/pnas.1921027117, (https://www.pnas.org/doi/abs/10.1073/pnas.1921027117) (2020).

47. K. J. Blumer, J. E. Reneke, J. Thorner, The STE2 gene product is the ligandbinding compo nent of the alphafactor receptor of Saccharomyces cerevisiae. Journal of Biological Chemistry 263, 10836–10842, ISSN: 00219258, DOI https://doi.org/10.1016/S0021-9258(18)38046-3, (https://www.sciencedirect.com/science/article/pii/S0021925818380463) (1988).

48. L. M. Leavitt, C. R. Macaluso, K. S. Kim, N. P. Martin, M. E. Dumont, Dominant negative mutations in the αfactor receptor, a G proteincoupled receptor encoded by the STE2 gene of the yeast Saccharomyces cerevisiae. Molecular and General Genetics MGG 261, 917–932, ISSN: 1432-1874, DOI 10.1007/s004380051039, (https://doi.org/10.1007/s004380051039) (Aug. 1999).

49. A. ColmanLerner et al., Regulated celltocell variation in a cellfate decision system. Nature 437, 699–706, ISSN: 1476-4687, DOI 10.1038/nature03998, (https://doi.org/10.1038/nature03998) (Sept. 2005).

50. E. Batchelor, M. Goulian, Robustness and the cycle of phosphorylation and dephosphoryla tion in a twocomponent regulatory system. Proceedings of the National Academy of Sciences 100, 691–696, DOI 10.1073/pnas.0234782100, eprint: https://www.pnas.org/doi/pdf/10.1073/pnas.0234782100, (https://www.pnas.org/doi/abs/10.1073/pnas.0234782100) (2003).

51. G. Shinar, M. Feinberg, Structural Sources of Robustness in Biochemical Reaction Networks. Science 327, 1389–1391, DOI 10.1126/science.1183372, eprint: https://www.science.org/doi/pdf/10.1126/science.1183372, (https://www.science.org/doi/abs/10.1126/science.1183372) (2010).

